# Genome editing of an African elite rice variety confers resistance against endemic and emerging *Xanthomonas oryzae* pv. *oryzae* strains

**DOI:** 10.1101/2022.11.20.517251

**Authors:** Van Schepler-Luu, Coline Sciallano, Melissa Stiebner, Chonghui Ji, Gabriel Boulard, Amadou Diallo, Florence Auguy, Si Nian Char, Yugander Arra, Kyrylo Schenstnyi, Marcel Buchholzer, Eliza P.I. Loo, Atugonza L. Bilaro, David Lihepanyama, Mohammed Mkuya, Rosemary Murori, Ricardo Oliva, Sebastien Cunnac, Bing Yang, Boris Szurek, Wolf B. Frommer

**Affiliations:** Institute for Molecular Physiology, Heinrich Heine University Düsseldorf, Düsseldorf, Germany; Division of Plant Science and Technology, Bond Life Sciences Center, University of Missouri, Columbia, MO 65211, USA; Plant Health Institute of Montpellier (PHIM), Université Montpellier, IRD, CIRAD, INRAE, Institut Agro, Montpellier, France; Tanzania Agricultural Research Institute (TARI)-Uyole Centre, PO Box 400 Mbeya, Tanzania; International Rice Research Institute, Eastern and Southern Africa Region (IRRI ESA), BOX 30709 00100, Nairobi, Kenya; International Rice Research Institute (IRRI), Africa Regional Office, Nairobi, Kenya; International Rice Research Institute, Pili Drive, Los Baños, Laguna 4031, Philippines; World Vegetable Center, Shanhua, Tainan, Taiwan 74151; Donald Danforth Plant Science Center, St. Louis, MO 63132, USA; Institute for Transformative Biomolecules, ITbM, Nagoya University, Nagoya, Japan

**Author notes:** Co-first authors, equal contribution (Van Schepler-Luu, Coline Sciallano, Melissa Stiebner, Chonghui Ji).

## Abstract

Bacterial leaf blight (BB) of rice, caused by *Xanthomonas oryzae* pv. *oryzae* (*Xoo*), threatens global food security and the livelihood of small-scale rice producers. Analyses of *Xoo* collections from Asia, Africa and the Americas demonstrated surprising continental segregation, despite robust global rice trade. Here, we report unprecedented BB outbreaks in Tanzania. The causative strains, unlike endemic *Xoo*, carry Asian-type TAL effectors targeting the sucrose transporter *SWEET11a* and suppressing *Xa1*. Phylogenomics clustered these strains with *Xoo* strains from China. African rice varieties do not carry suitable resistance genes. To protect African rice production against this emerging threat, we developed a hybrid CRISPR-Cas9/Cpf1 system to edit six TALe-binding elements in three *SWEET* promoters of the East African elite variety Komboka. The edited lines show broad-spectrum resistance against Asian and African strains of *Xoo*, including strains recently discovered in Tanzania. This strategy could help to protect global rice crops from BB pandemics.

## Introduction

Rice is one of the most important staple foods for developing countries in Asia and Africa (Odongo et al., 2021). African consumers increasingly replace the traditional staples of sorghum, millet and maize with rice. Today, African farmers, 90% of which are small-scale food producers with <1 ha of land (Pandey et al., 2010), produce ~60% of the local rice demand, but this demand will increase with the expected doubling of the population until 2050. Productivity is often hampered by the diseases Bacterial Leaf Blight (BB), Rice Yellow Mottle Virus (RYMV), and Rice Blast (RB)(Jiang et al., 2020; Longue et al., 2018; Mutiga et al., 2021). Breeding high-yielding varieties that are resistant to these diseases will be an important factor supporting food security in Africa. BB, caused by the bacterium *Xanthomonas oryzae* pv. *oryzae (Xoo)*, is a devastating rice disease in many rice-growing countries. Resistance (*R*) genes for BB have been identified and used extensively to develop resistant varieties, however resistance based on single *R* genes may be rapidly overcome by new strains (Vera Cruz et al., 2000). Regular monitoring of the virulence of current *Xoo* strains on a collection of rice tester lines carrying single *R* genes has provided guidance for stacking different *R* genes to obtain broad-spectrum resistance in Asian rice varieties (Arra et al., 2018, 2017).

In Africa, BB was first reported in Mali in 1979 and later in several other West and Central African countries (Buddenhagen et al., 1979; Verdier et al., 2012b). Recently, East African countries like Tanzania and Uganda reported BB epidemics (Duku et al., 2016). While the information on the spread and severity of BB in Africa is scarce, BB is not yet considered a major threat in Africa, but has established. However, climate change affects disease spread, leading to projections that the damage will become more severe in the future (Amos, 2013; Séré et al., 2005; Verdier et al., 2012b). Other factors, such as population growth and increasing yield losses due to pathogens, as well as the distinct *Xoo* populations in Africa, make it essential to generate broad-spectrum and durable resistance in local varieties specifically for Africa.

Key aspects of the mechanisms underlying BB disease have been elucidated and provide an efficient roadmap to breeding resistance. *Xoo* secretes a series of Transcription Activator-Like effectors (TALes) into rice xylem parenchyma cells via a type III secretion system. Once inside the host cell, the TALes translocate to the nucleus and bind to effector binding elements (EBEs) in the promoters of specific host genes via a unique domain of tandemly arranged 34 amino-acid long repeats, triggering the ectopic induction of the target genes (Richter et al., 2014). *Xoo* TALes induce one or several host SWEET sucrose uniporter-genes (*SWEET11a, 13* and *14*), presumably causing sucrose release into the apoplasm where bacteria reside. Note that due to the discovery of a previously misannotated sucrose transporting paralog, SWEET11a was renamed (formerly SWEET11)(Wu et al., 2022). Sucrose is consumed by the bacteria, resulting in effective reproduction (Sadoine et al., 2021). Allelic EBE variants that are not recognized by TALes function as recessive gene-for-gene resistance genes. To date, six EBEs in *SWEET* promoters are known for the following TALes: PthXo1 in *SWEET11a*, PthXo2 variants in *SWEET13*, and TalC, PthXo3, AvrXa7 and TalF in *SWEET14* (Eom et al., 2019; Oliva et al., 2019). Notably, Asian and African *Xoo* strains are phylogenetically distinct and use different TALes to target *SWEETs* (Oliva et al., 2019; Tran et al., 2018). African *Xoo* strains exclusively use TalC and TalF, which both target *SWEET14*, while Asian strains encode PthXo1, PthXo2 (variants A, B, C), PthXo3 and AvrXa7, which target *SWEET11a, 13* and *14*. American *Xoo* strains lack TALes and hence are poorly virulent (Verdier et al., 2012a). Based on this information, broad-spectrum resistance was introduced into the *Oryza sativa* ssp. *japonica* variety Kitaake and the spp. *indica* varieties IR64 and Ciherang-Sub1 by genome editing of the EBEs in all three *SWEET* genes (Eom et al., 2019; Oliva et al., 2019). The edited lines may prove to be valuable breeding materials, giving rise to lines that will benefit small-scale producers in Asia. To prepare for emerging strains with novel virulence mechanisms, the diagnostic SWEET^R^ kit was developed, enabling analyses of BB in the field and identification of suitable resistance strategies, such as which *R*-gene combinations to deploy in the next season (Eom et al., 2019).

Recent disease surveys in Tanzania, described here, have identified a BB outbreak in 2019 that has spread in subsequent years. The causative strains have features that distinguish them from other African *Xoo* strains and have virulence properties that make them a major threat to African rice production. Recently, Kenya and Nigeria have exempted SDN-1 genome-edited crops generated using site-directed nuclease without a transgene (SDN-1) from GMO regulations (Buchholzer and Frommer, 2022). This exemption provides a regulatory framework for the introduction of genome-edited, BB-resistant rice into Africa. To provide small-scale rice producers with BB-resistant rice cultivars, we used an efficient transformation protocol (Luu et al., 2020) to introduce BB resistance into a rice cultivar popular in East African countries. Komboka (IRO5N-221) is a high-yielding, semi-aromatic rice variety jointly developed by IRRI (International Rice Research Institute and KALRO (Kenya Agricultural & Livestock Research Organization; <u>BBSRC Varieties Kenya.pdf</u>) that grows to an average height of 110 cm and has a yield potential of ~7 tons/ha, almost twice that of Basmati, a popular variety planted in Kenya and Rwanda (<u>The Star 2020</u>). Komboka plants mature in ~ 115 days, respond well to fertilization, and are suitable for irrigated lowland cultivation in Africa (Kitilu et al., 2019). Komboka had been classified as moderately resistant to BB, RYMV, and RB, however, detailed resistance profiles are not available. In this work, Komboka BB-resistance genes were analyzed, and a combination of Cas9, Cpf1 and multiplexed gRNAs was used to edit all known EBEs in the *SWEET* genes, resulting in broad-spectrum resistance to not only previously known African and Asian strains, but also the newly introduced *Xoo* strains from the outbreak in Tanzania.

## Results

### Identification of an unprecedented disease outbreak in Tanzania

There is little information on the population genetics and dynamics of *Xoo* strains circulating in Africa. While pathogenic strains have sporadically been isolated from rice plants and characterized, there is no systematic analysis of BB across Africa. Moreover, there are many diverse rice varieties used in Africa, with no general survey of acreage growing different varieties. These factors combine to make it challenging to breed durably resistant lines. In addition, numerous international partnerships have introduced varieties and cultivation techniques to major rice production areas in different countries of Africa. For instance, in 1975, one of the first irrigated perimeters with rice research labs and a training farm were developed in Dakawa, a major rice production zone in Tanzania, as a result of a multi-year partnership with North Korea (van der Tycho and Yonho, 2022). National (NAFCO) and international institutions (USAID, Cornell University) remain active in Dakawa to build irrigation schemes and implement *The System of Rice Intensification* (SRI; http://sri.ciifad.cornell.edu/countries/tanzania/index.html#summary). In 2011, a collaboration with China resulted in 50 ha of irrigated fields managed by local farmers. There is also a Technology Demonstration Center that trains local farmers and technicians and grows examples of Chinese improved varieties, including hybrid rice (Makundi, 2017).

The first observations of BB in Africa were made in Mali in 1979 (Buddenhagen et al., 1979). Since then, the disease has been reported mostly in West Africa, including Senegal, Benin, Burkina Faso, Ivory Coast, Mali and Niger (Afolabi et al., 2016; Gonzalez et al., 2007; Tall et al., 2020; Tekete et al., 2020). More recently, BB was also reported in the East African countries Uganda and Tanzania (Duku et al., 2016; Oliva et al., 2019). To systematically analyze these newly isolated strains and to compare them to the broader African *Xoo* landscape, 833 strains from rice fields sampled in nine African countries between 2003 and 2021 were collected and characterized using molecular diagnostic as well as pathogenicity assays (Supplementary Table 1)(Afolabi et al., 2016; Gonzalez et al., 2007; Tall et al., 2020; Tekete et al., 2020). During this study, two unprecedented outbreaks were identified, one in 2019 in Dakawa and another in 2022 in Lukenge (Supplementary Table 2). These sites are located 60 km apart in the Morogoro region of Tanzania, an area that for decades has been a center of partnership initiatives aiming at increasing rice production (Makundi, 2017). We performed disease surveys in this area in 2019 and 2021 and identified two outbreaks on TXD 306 (SARO-5), a rice variety popular in irrigated ecologies in Tanzania. More recently BB was identified on Komboka. Leaves with typical BB symptoms were processed, and seven strains from Dakawa and 106 from Lukenge were isolated and validated as *Xoo* with diagnostic primers (Lang et al., 2010). Notably, surveys performed in subsequent years showed increased severity and spread (Supplementary Table 2).

To identify the *R* genes best suited for BB control, 21 strains that represent the diverse geographical origins of our collection were inoculated on six Near Isogenic Lines (NILs) of rice that were harboring either *Xa1, Xa3, Xa4, xa5, Xa7* or *Xa23* and that had been reported as efficient *R* genes against some African *Xoo* strains (Table 1, Supplementary Table 1)(Tekete et al., 2020). The test panel included three strains, namely CIX4457, CIX4458 and CIX4462, that were isolated from the recently detected outbreak in Dakawa, Tanzania. NILs harboring *Xa1, Xa23, xa5* or *Xa4* were resistant to most African strains, while CIX4457, CIX4458 and CIX4462 were virulent on all NILs tested, highlighting an urgent need for a new solution to protect rice against BB outbreaks.

### Resistance of Komboka to endemic African *Xoo* but not to new Tanzanian strains

Komboka, an emerging elite variety released in some countries in East Africa (Tanzania, Kenya, Uganda and Burundi), has been described as moderately resistant to *Xoo*, yet the nature of its resistance has not been elucidated. Using the African *Xoo* diversity panel, we found that most African strains were avirulent in Komboka, except for the newly isolated strains from Tanzania (Fig. 1). By comparison, all strains were highly virulent on the susceptible variety Azucena (Figure 1-figure supplement 1, Table 1). Since *Xa1*, *Xa4, xa5* and *Xa23* confer resistance against *Xoo* (Table 1), we investigated potential *R* genes in Komboka. Mining of the IRRI QTL database (https://rbi.irri.org/resources-and-tools/qtl-profiles) revealed that Komboka contains genetic markers linked to *Xa4* but not *xa5* or *Xa23*, while there was no information on *Xa1* in the database (Figure 1-figure supplement 2A). We confirmed the presence of *Xa4* in Komboka by tracing an *Xa4*-associated marker and by sequencing (Figure 1-figure supplement 2B, C). Consistent with the broad-spectrum resistance of *Xa1* against endemic African *Xoo* strains, we found that Komboka carries a dominant *Xa1* allele that is highly similar to *Xa45*(t) (Figure 1-figure supplement 2D, E). The combination of *Xa4* and *Xa45(t)* in Komboka can explain the observed resistance to most African *Xoo* strains. Apparently, the two *R* genes do not protect against the strains recently isolated from the Dakawa outbreaks.

**Figure 1.**
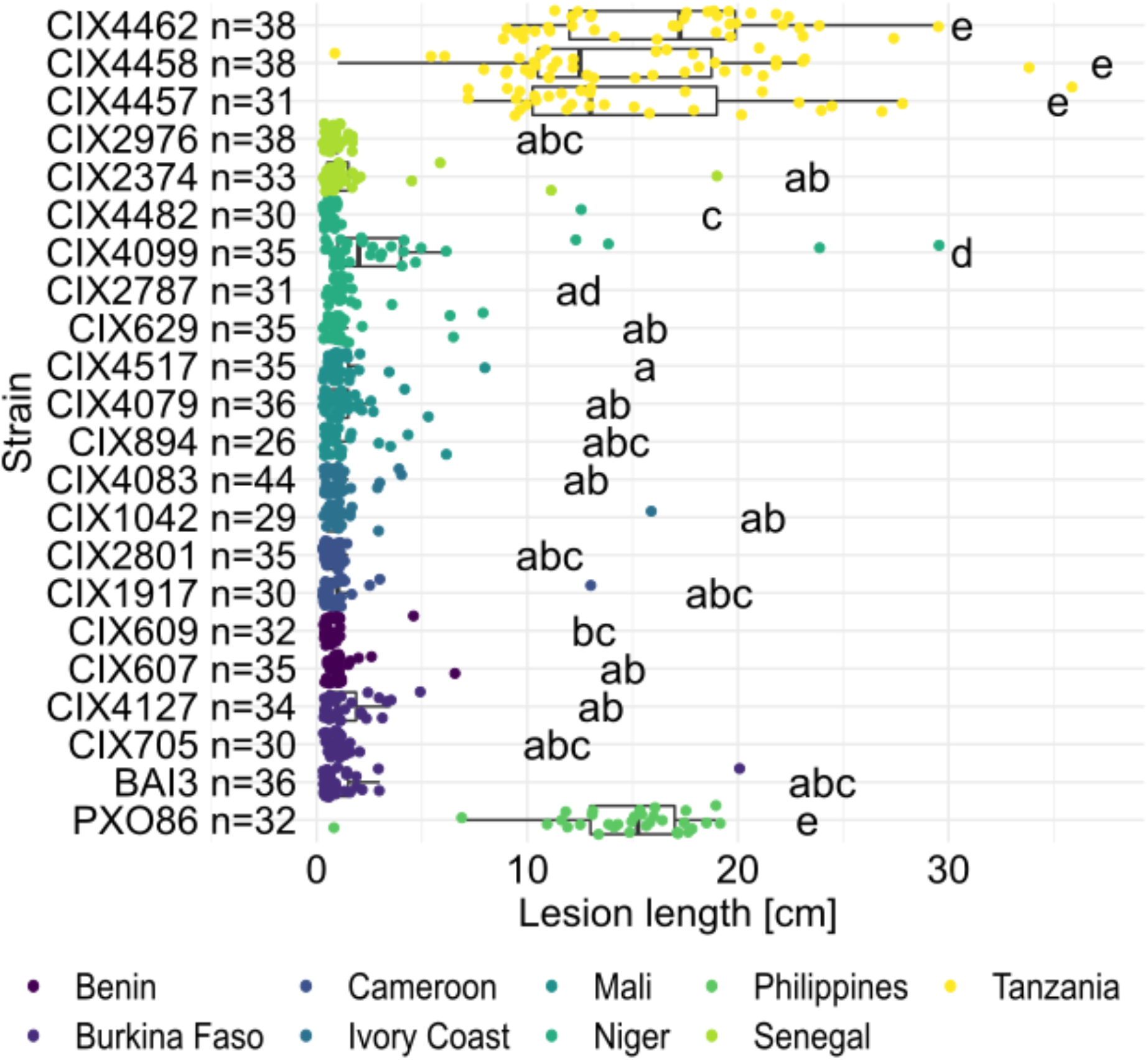
Resistance spectrum of wild-type Komboka rice against African *Xoo* strains. Leaf-clip inoculation of wild-type Komboka rice plants with a panel of 21 *Xoo* strains originating from 8 African countries along with Asian reference strain PXO86 from the Philippines. Lesion length (in cm) was measured 14 days after *Xoo* inoculation. Data from three independent experiments are represented. Letters in the boxplot represent significant differences.

**Table 1:** Efficiency of resistance genes in IRBB rice lines towards a diversity panel of 21 African *Xoo* strains. Qualitative scores of the lesion lengths produced upon leaf-clip inoculation of a diversity panel consisting of 21 *Xanthomonas oryzae* pv. *oryzae* strains from West and East Africa on IRBB near-isogenic-lines (NILs) carrying different *Xanthomonas* resistance genes **(*Xa*)** and on the variety Azucena used as susceptible control (IRBB1: *Xa1;* IRBB3: *Xa3;* IRBB4: *Xa4*; IRBB5: *xa5;* IRBB7: *Xa7;* IRBB23: *Xa23;* IR24: susceptible control). Resistance or susceptibility of rice plants to *Xoo* was determined by lesion length measured 14 days after inoculation: Resistant (R) < 5 cm; moderately resistant (MR) = 5 to 10 cm; moderately susceptible (MS) = 10 to 15 cm; and susceptible (S) >15 cm.

| CIX Number | Strain Name | Country | Azucena | IR24 | IRBB1 | IRBB3 | IRBB4 | IRBB5 | IRBB7 | IRBB23 |
| --- | --- | --- | --- | --- | --- | --- | --- | --- | --- | --- |
| 607 | Nati Park | Benin | S | R | R | R | R | R | R | R |
| 705 | Karfiguela 13 | Burkina Faso | S | R | R | R | R | R | R | R |
| 4482 | N2-4 | Niger | S | R | R | R | R | R | R | R |
| 609 | Tanguieta3 | Benin | S | MR | R | MR | R | R | MR | R |
| 629 | Toula20 | Niger | S | MR | R | MR | R | R | MR | R |
| 4127 | BAI250 | Burkina Faso | S | MR | R | MR | MR | R | R | R |
| 2374 | S62-2-22 | Senegal | S | MR | R | MS | R | R | MR | R |
| 4083 | CII-1 | Ivory Coast | S | MS | MR | MS | R | R | MR | R |
| 1042 | CII-2 | Ivory Coast | S | MS | MR | MS | MR | R | MR | R |
| 2787 | NAI9 | Niger | S | MS | MR | MS | MR | MR | MS | R |
| 1917 | AXO1947 | Cameroon | S | S | R | S | R | R | MS | R |
| 894 | MAI145 | Mali | S | S | R | S | MR | MS | S | R |
| 4092 | BAI3 | Burkina Faso | S | MS | MR | S | MR | MR | MS | R |
| 2801 | CFBP1948 | Cameroon | S | MS | MR | S | MR | MR | S | R |
| 4099 | NAI5 | Niger | S | S | MR | S | MR | MR | S | R |
| 4079 | MAI73 | Mali | S | S | R | S | MS | MR | S | R |
| 2976 | S82-4-3 | Senegal | S | MS | MR | S | MS | MR | MS | R |
| 4517 | MAI132 | Mali | S | S | R | S | S | MS | S | R |
| 4457 | Tz1 | Tanzania | S | S | S | S | S | S | S | S |
| 4458 | Tz2 | Tanzania | S | S | S | S | S | S | S | S |
| 4462 | Tz6 | Tanzania | S | S | S | S | S | S | S | S |

### Tanzanian *Xoo* strain cluster with Asian *Xoo* via whole genome SNP-based phylogeny

To identify the mechanisms underlying the virulence of the Tanzanian strains, three representative strains: CIX4462 (Tz6) collected from Dakawa in 2019 and CIX4506 (F2SP1-1c) and CIX4509 (F4SP1-1a), collected from Lukenge in 2021, were subjected to whole-genome sequencing and SNP-based phylogenetic analyses (Supplementary Table 3). Notably, CIX4457, CIX4458 and CIX4462 clustered with Asian *Xoo* isolates (Fig. 2A). In contrast, the Tanzanian strains T19, Dak16 and Xoo3-1, which had been collected in Dakawa before the 2019 outbreak, grouped with the endemic African lineage, consistent with previous reports (Oliva et al., 2019). CIX4462, CIX4506 and CIX4509 are most closely related to strains from Yunnan province, China (Fig. 2B) and carry *iTALe* genes similar to *tal3a* (Fig. 2C). *iTALe* genes encode truncated TALes known to suppress *Xa1* resistance, but had so far only been reported in Asian *Xoo* isolates. In addition, the three strains contain a TALe (named PthXo1B) highly similar to the Asian PthXo1, with the addition of two RVD domains at the N-terminus (Fig. 2D; Supplementary Data 1). EBE prediction indicated that PthXo1B can bind to the *SWEET11a* promoter at a site overlapping with the EBE targeted by PthXo1 (Fig. 2D). Importantly, *SWEET11a* knock-out mutants from the diagnostic SWEET^R^ kit were resistant to CIX4457 and CIX4505 from Tanzania, whereas *sweet13* or *sweet14* knock*-*out mutants remained susceptible (Figure 3A). In accordance, *SWEET11a* was induced in Kitaake leaves when inoculated with the Tanzanian strains CIX4457 and CIX4505 (Figure 3B). Together, our data showed that representative Tanzanian strains collected in 2019-2021 are phylogenetically related to Asian *Xoo* strains and have both *iTALe* and *PthXo1* homologs unique to Asian *Xoo* isolates (Oliva et al., 2019).

**Figure 2.**
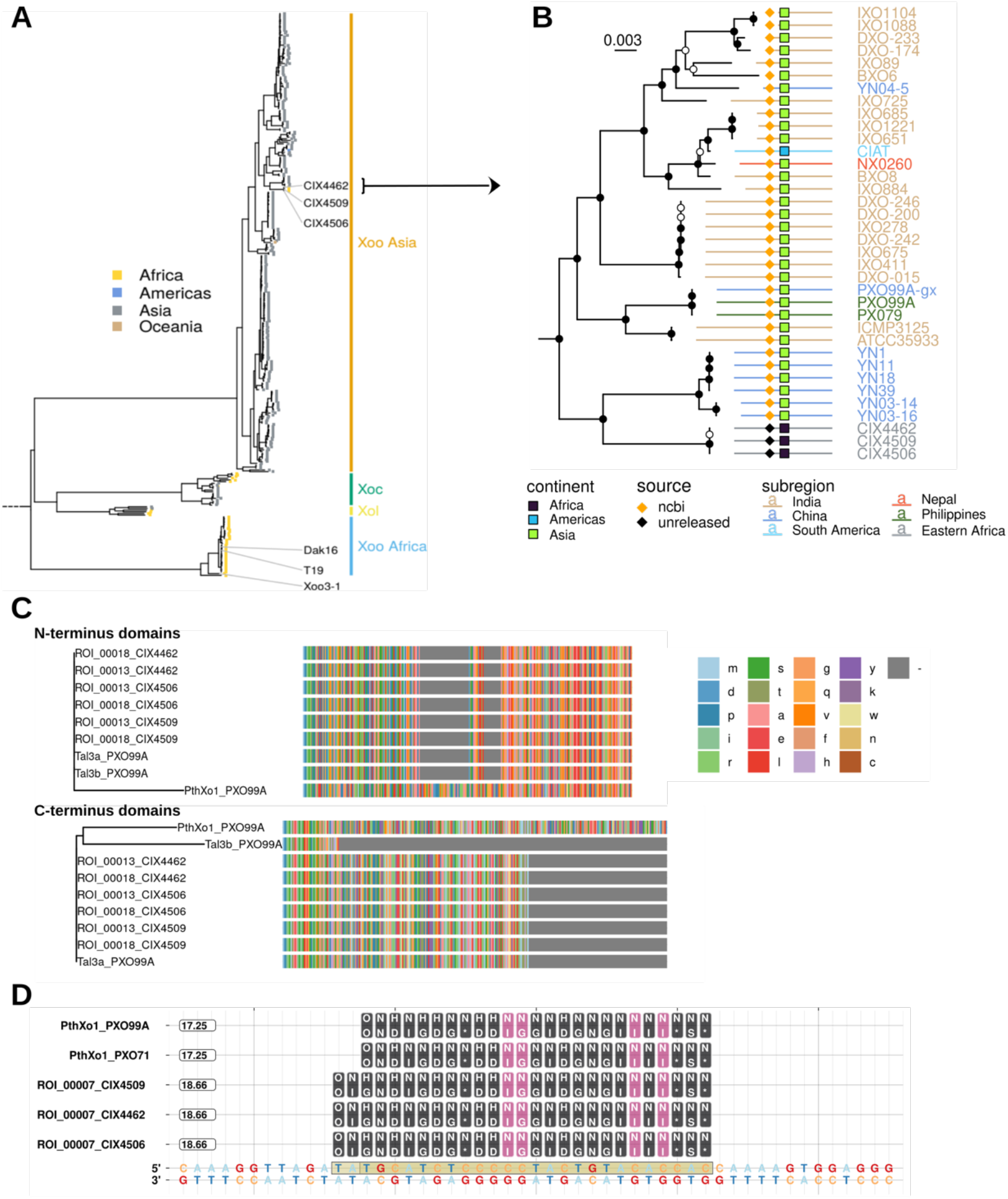
Analysis of the genomes of Tanzanian *Xoo* isolates. **A.**Core genome *Xanthomonas oryzae* phylogenetic tree. Only the names of Tanzanian isolates are indicated. Abbreviations: *Xoo*, *X. oryzae* pv. *oryzae; Xoc, X. oryzae* pv. *oryzicola; Xol, X. oryzae* pv.*leersiae*.**B.**Close-up view of a portion of the tree in A, focusing on the newly isolated Tanzanian strains and neighboring clades. Black-filled nodes have a bootstrap support value equal to or above 80%. The scale bar reflects branch length in mean number of nucleotide substitutions per site. **C.**Miniature multiple alignment of the terminal domain sequences of PXO99A iTALes and the putative iTALes from the three newly isolated Tanzanian strains. The PXO99A PthXo1 domains act as ‘standard’ TALe references, with lowercase letters representing amino acids. Gaps are colored in gray. **D.**Talvez EBE predictions on the *SWEET11a* promoter. Repeat Variable Di-residue (RVD) sequences (rounded boxes) are aligned along their predicted matching nucleotide along the promoter sequence (the lowest row). Black-filled RVDs match their target nucleotide in the Talvez RVD-nucleotide association matrix with the best possible score for this RVD. Those in violet match with an intermediate score. Values in the rounded boxes near the TALe names correspond to Talvez prediction scores.

**Figure 3.**
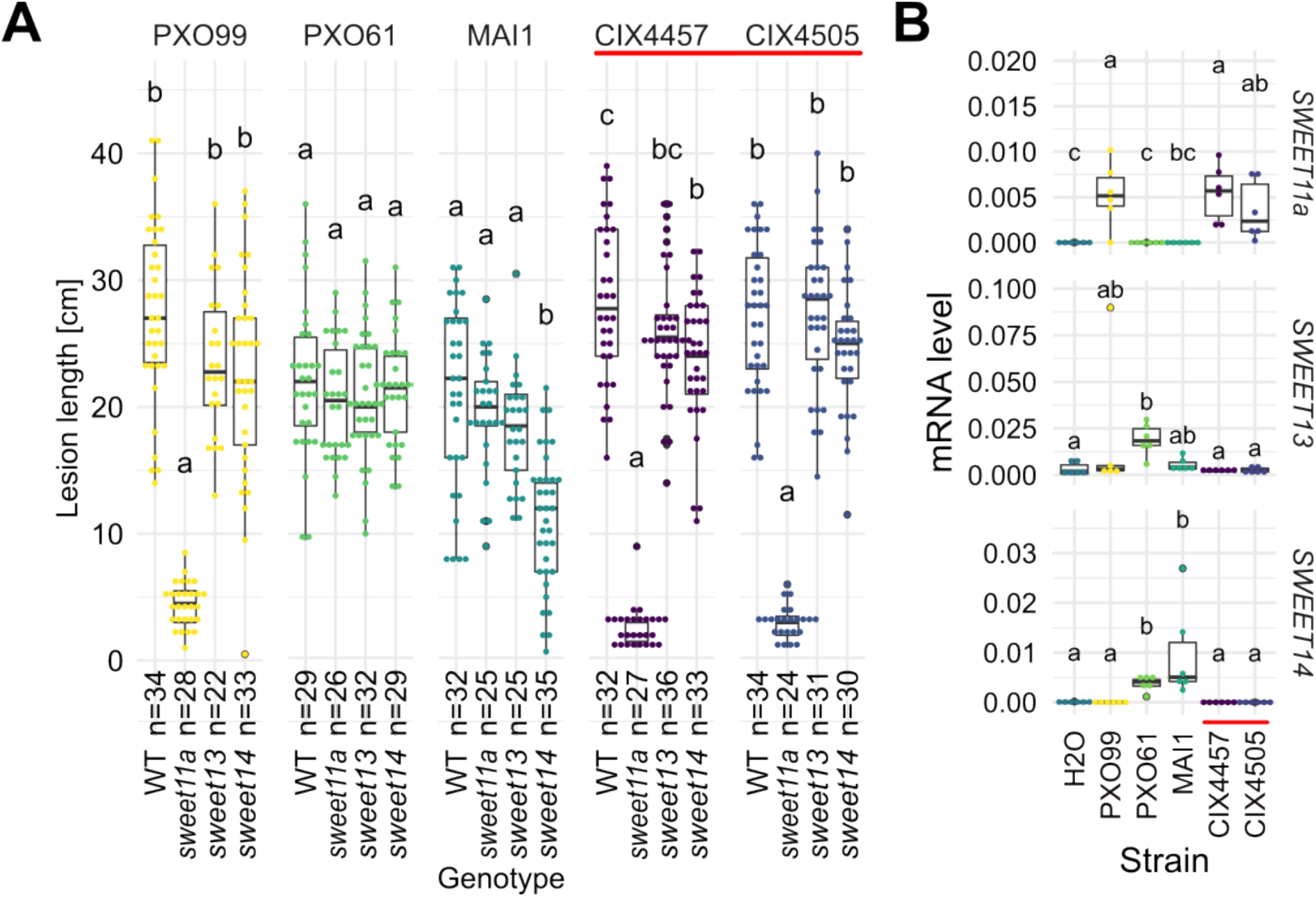
Tanzanian *Xoo* strains rely on the induction of *SWEET11a* to cause disease. **A.**Lesion lengths (in cm) were measured 14 days after leaf-clipping inoculation of Kitaake individual *sweet* knock out lines (sweet11a, sweet13 and sweet14) in the cultivar Kitaake (Eom et al., 2019) with PXO99 (PthXo1), PXO61 (PthXo2B/PthXo3), MAI1 (TalC/TalF), and Tanzanian strains (CIX4457 and 4505) from the recent outbreaks (highlighted by a red bar). Results from two independent experiments are represented. **B.**Relative mRNA levels (2^-ΔCt^) of *SWEET11a, SWEET13* and *SWEET14* in wild-type Kitaake upon infection by PXO99 (PthXo1), BAI3 (TalC), and Tanzanian *pthXo1B* dependent *Xoo* strains. Samples were collected 48 h post infiltration. Data from three independent experiments were pooled and are represented together. All Ct values were normalized by rice *EF1α* elongation factor (△Ct).

### Editing *SWEET* promoter sequences in Komboka to obtain resistance to all African strains

As a prerequisite for editing *SWEET* promoters in Komboka, the promoter regions of *SWEET11a, 13* and *14* were sequenced and shown to contain EBEs for PthXo1, PthXo2A, PthXo3, AvrXa7, TalC and TalF (Figure 4-figure supplement 1, Supplementary Table 4, Supplementary Data 2). To protect Komboka against endemic strains from Africa as well as introduced Asian strains, all six known EBEs in the promoters of *SWEET11a, 13* and *14* were edited using a hybrid Cas9/Cpf1 editing system. Due to blunt cleavage, Cas9 preferentially produces SNPs, while Cpf1 (Cas12) produces staggered cuts, creating predominantly small deletions. We hypothesized that small deletions would produce more robust resistance. Due to the PAM requirement, it was not possible to design sgRNAs for Cpf1 at all EBEs; therefore, Cpf1 was combined with Cas9. Two CRISPR/Cpf1 gRNAs (cXo1 and cXo2) were designed to target PthXo1 and PthXo2A EBEs within the promoters of *SWEET11a* and *SWEET13*,respectively (Figure 4-figure supplement 1, 2). Since the EBEs for AvrXa7, PthXo3 and TalF in the promoter of *SWEET14* are overlapping, one CRISPR-Cpf1 gRNA (cTalF) was designed to target the overlapping region of all three EBEs. Because no TTTV PAM sequence was available near the EBE for TalC, a Cas9 gRNA (gTalC) was designed to target TalC EBE. From a first round of transformation, six representative T2 lines were tested for resistance using six representative *Xoo* strains: ME2 (PXO99A mutant deficient in PthXo1), PXO99A (containing PthXo1), PXO61 (PthXo2B, PthXo3), PXO86 (AvrXa7), MAI1 (TalC, TalF) and BAI3 (TalC) (Fig. 4A, Figure 4-figure supplement 3, Supplementary Table 4, 5, 6). All six lines from this first round of transformation were resistant to the tested strains. One line (1.5_19) was fully resistant to all strains, while five lines were fully resistant to five strains but only moderately resistant to PXO86. This difference in resistance is most likely explained by the presence of a 4-bp deletion in the EBE for AvrXa7, while line 1.5_19 carried the same 4-bp deletion plus an additional base pair substitution (G/T) (Fig. 4B). Since the clipping assays used very high bacterial titers, it is generally assumed that moderate resistance will be sufficient for effective resistance in field conditions. However, TALes were reported to have less specific nucleotide binding at the 3’-end, hence the G/T substitution in line 1.5_19 could potentially be overcome (Richter et al., 2014). To obtain more robust resistance to a larger population of *Xoo*, a second round of transformation was carried out. Two T2 lines (14_19 and 14_65), which contained deletions in all EBEs, including 11-bp and 5-bp deletions in the AvrXa7 EBEs, respectively, were resistant to all tested strains including PXO86 (Fig. 5). Lines 1.5_19 and 1.2_40 were also resistant to CIX4457 and CIX4505, the new isolates from Tanzania, consistent with the 12- and 9-bp deletions in the predicted PthXo1 EBE, respectively (Fig. 6). As one may have predicted, induction of *SWEET11a* by CIX4457 and CIX4505 was abolished in lines 1.2_40 and 1.5_19 (Figure 6-figure supplement 1). Taken together, the edited Komboka lines are fully resistant to Asian and African *Xoo* strains, including the emerging population from Dakawa and Lukenge in Tanzania.

**Figure 4.**
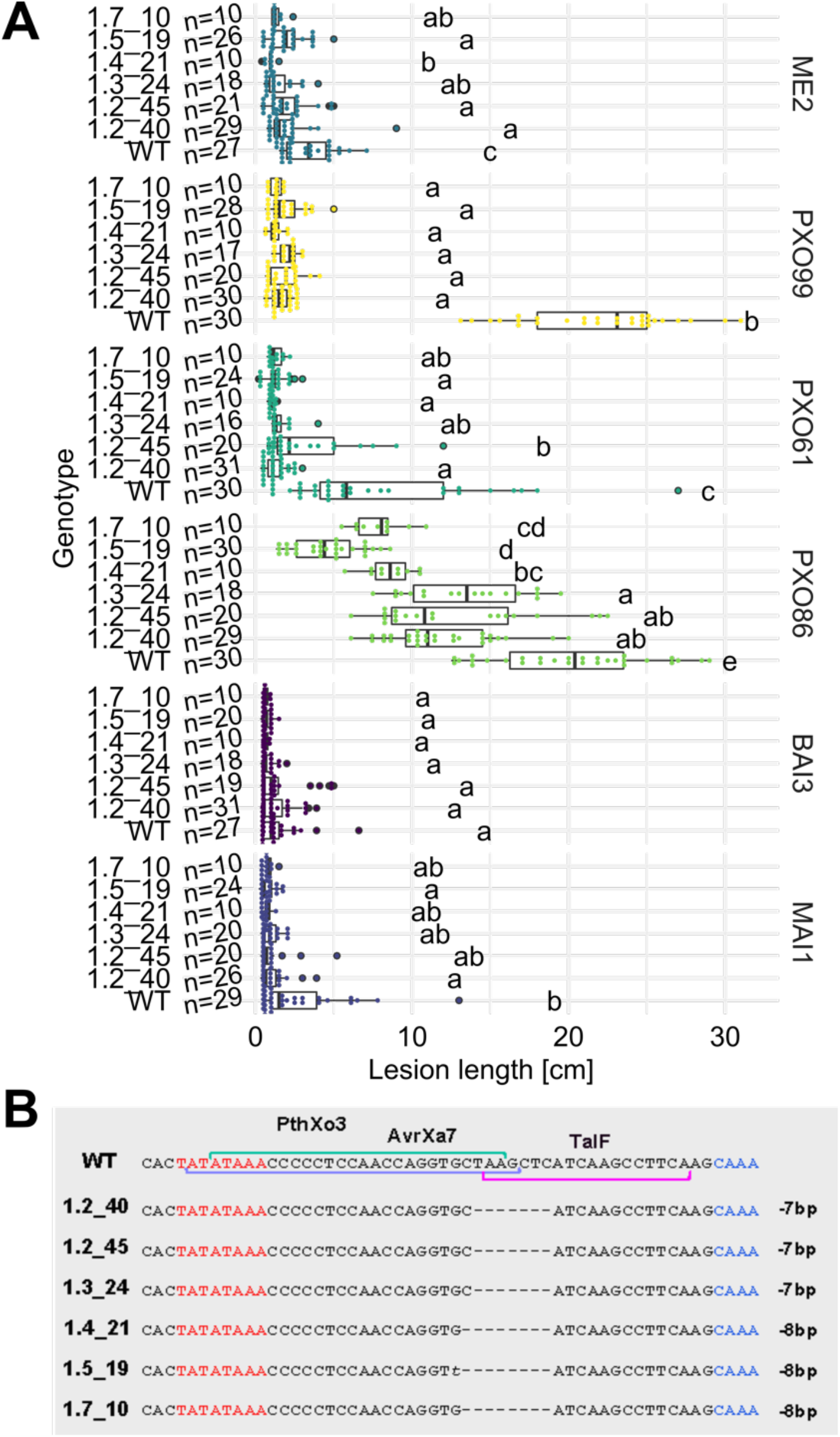
Resistance of EBE-edited Komboka lines against six representative *Xoo* strains. **A.**Reactions of WT and six edited Komboka lines to infection by *Xoo* strains (PXO99A, PXO61, PXO86, BAI3 and MAI1) harboring PthXo1, PthXo2 PthXo3, AvrXa7, TalC and/or TalF. ME2 is a PXO99A mutant strain with an inactivated *pthXo1* and served as a negative control. The Komboka lines were determined to be edited in all 6 EBE sites by sequencing. **B**. Details of the EBEs targeted by PthXo3, AvrXa7 and TalF and their mutations in six edited lines.

**Figure 5.**
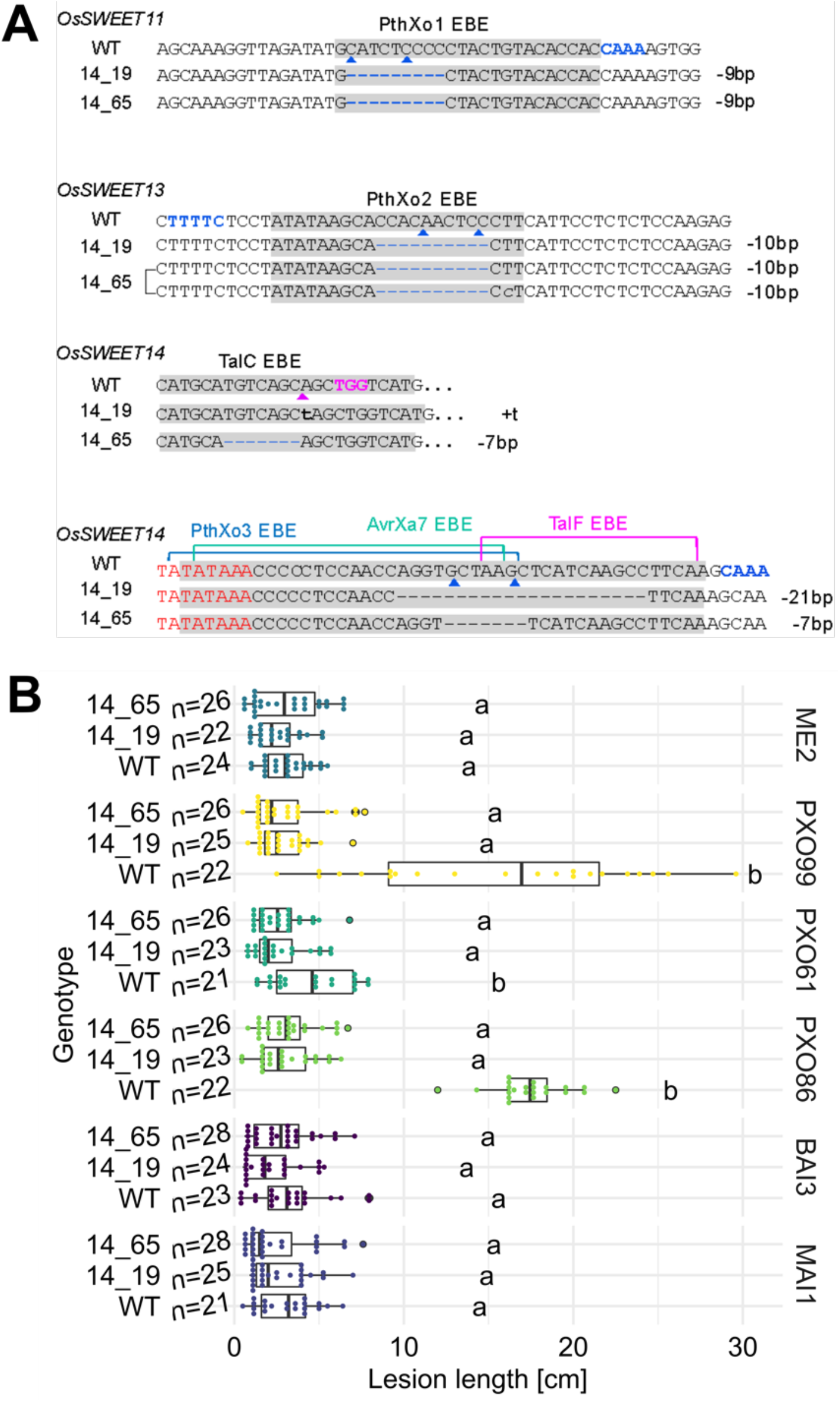
Genotypes and phenotypes of two EBE-edited lines generated in a second transformation experiment. **A.**Mutations at the EBEs for PthXo1, PthXo2, TalC, TalF, PthXo3, and AvrXa7 in two Komboka edited lines, 14_19 and 14_65. **B**. Reactions of wild-type Komboka and the two edited lines to the infections by PthXo1-, PthXo2-, PthXo3-, AvrXa7-, TalC-, and TalF-harboring *Xoo* strains (PXO99A, PXO61, PXO86, BAI3 and MAI1). ME2 is a PXO99A mutant that is inactivated in *pthXo1* and served as a negative control.

**Figure 6.**
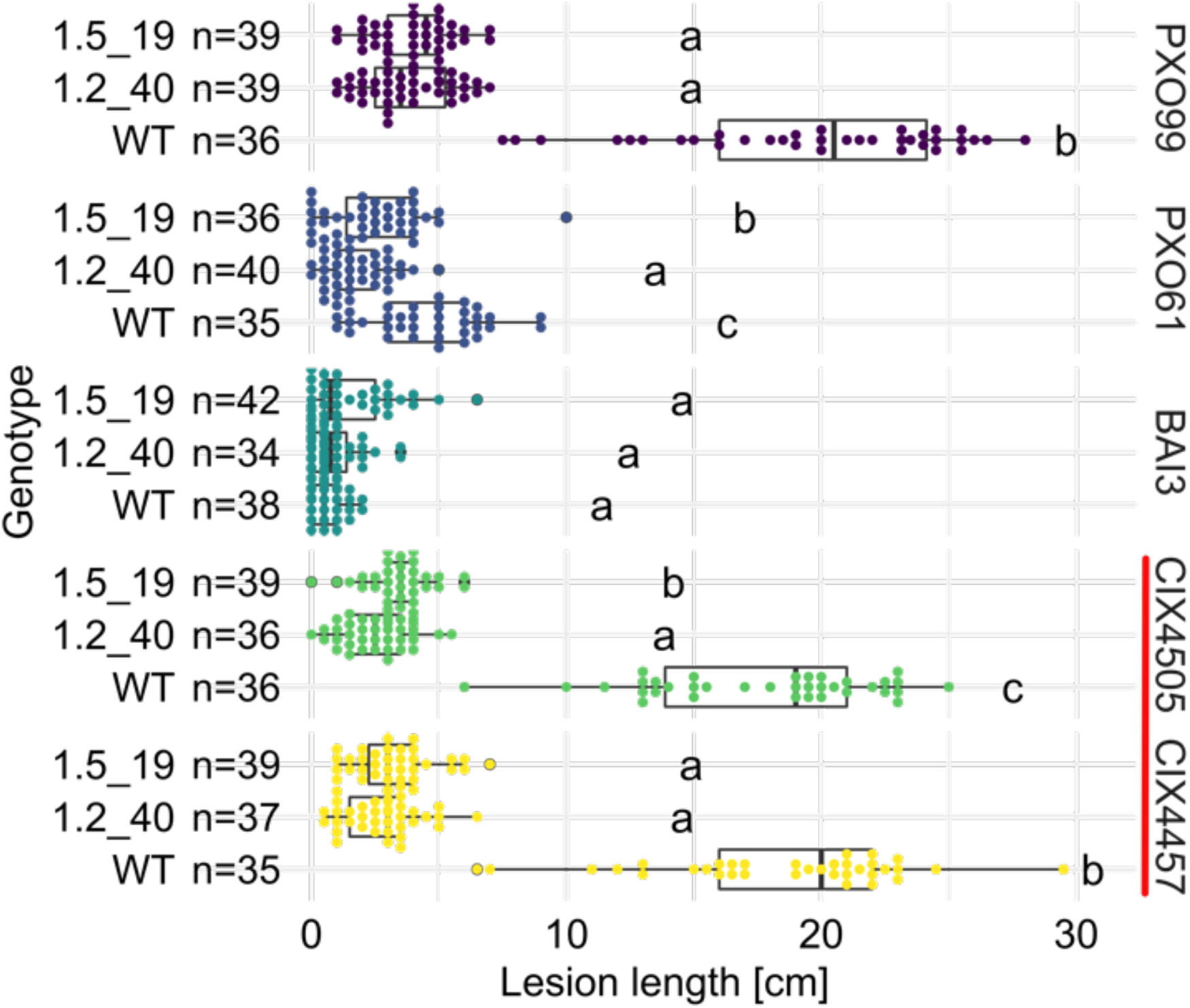
Komboka *SWEET* promoter edited lines are fully resistant against Tanzanian strains of *Xoo*. Lesion lengths (in cm) measured 14 days after leaf-clip inoculation of wild-type Komboka and the two multi-edited lines 1.2_40 and 1.5_19 with *Xoo* strains PXO99 (PthXo1), BAI3 (TalC), PXO61 (PthXo2 & PthXo3), and the newly isolated Tanzanian strains CIX4505 and CIX4457 (highlighted by a red bar). Four independent experiments are represented.

## Discussion

Here we report an outbreak of BB in East Africa. Through genetic and genomic characterization, this outbreak can be ascribed to a recent colonization event by *Xoo* strains that are most similar to Asian strains, specifically from Yunnan province.

Notably, the Asian and African strains clearly form distinct phylogenetical groups (Lang et al., 2019; Tran et al., 2018). This came as a surprise since monitoring had indicated very low BB incidence, and only three strains were isolated between 2011 and 2018 (Oliva et al., 2019). As shown previously, those strains cluster with other African *Xoo* strains using SNP-based phylogeny analyses and carry TalC as the major virulence TALe (Oliva et al., 2019). We cannot trace the introduction of the new strains, which may have occurred through either major storms or transfer of contaminated rice. *Xoo* is well-known to be transmitted via tropical storms, and transmission within Asian countries has been implied, for example in the case of PXO99A, which may have been introduced rather recently from Nepal or India to the Philippines. By contrast, rather effective continental isolation had been observed, and if the new Tanzanian strains would have been transmitted by storms from Asia, one may have expected a closer phylogenetic relationship to strains from India or Indonesia based on the geography (Oliva et al., 2019; Quibod et al., 2016; Tran et al., 2018). Due to the distinct evolutionary trace and the resulting absence of coevolution, it is likely that many African rice varieties do not contain suitable *R*-genes that could protect against Asian strains. Hallmark features of the new strains identified in Tanzania are iTALes and a PthXo1 homolog; both are absent from endemic African strains. African rice lines carrying *Xa1* may thus be resistant to African strains, but will be susceptible to Asian strains, including those recently introduced to Tanzania. Surveys over several years with preliminary data from 2022 indicate that the outbreak in Dakawa is gaining momentum regarding severity and spread, thus potentially becoming a threat to, at the least, East Africa but possibly to all of Africa by the increasing occurrence of heavy storms.

Our study, which so far is based on only a few strains representative of both outbreaks in Tanzania, shows that they are very close genetically, arguing in favor of a single introduction event. However, further analysis of a greater number of strains will be required to validate their origin and relationships. To gain insight into the spread of these emerging *Xoo* populations, pathogen monitoring was further conducted in Morogoro and surrounding regions in July 2022. Strikingly, over 600 leaf samples harboring typical BB symptoms were collected from 37 fields in six different regions in Tanzania. MLVA-typing and/or whole genome SNP analysis of the strains associated with these lesions will reveal the genetic structure of the *Xoo* population in Eastern Tanzania.

We had identified *SWEETs* as essential susceptibility loci for BB (Chen et al., 2012; Eom et al., 2019; Streubel et al., 2013). Virulence appears to be critically dependent on the activation of one of the sucrose-transporting SWEET uniporters in the rice genome (Oliva et al., 2019). Another relevant difference between Asian and African strains is the fully distinct set of TALes used to target SWEETs (Oliva et al., 2019). It is thus likely that, due to the absence of coevolution, African rice varieties contain EBEs in *SWEET* promoters that can be targeted by Asian TALes.

Analysis of Komboka, a high yielding semi-aromatic rice line released in Kenya and Tanzania, shows that it is resistant to endemic African strains due to *Xa1* and *Xa4*. But Komboka is susceptible to Asian strains like PXO99A, which carries both iTALes and PthXo1 that suppress Xa1 and induce *SWEET11a*,respectively. Indeed, we found Komboka samples with BB symptoms in another district in Tanzania (Mikindani, Mtwara, ~400 km from Dakawa) in 2022. Characterization of the collected leaves is ongoing. *Xa1*-mediated resistance has been broken by 95% of the Asian *Xoo* strains (Ji et al., 2020), while *Xa4*-mediated resistance was also overcome by the new East African *Xoo* strains as well as many Asian strains reported previously (Quibod et al., 2016). Thus,*Xa1* and *Xa4* lack durability due to shifts in the *Xoo* population or the emergence or introduction of new strains. Thus new lines with broadspectrum resistance against both Asian and African strains, including the strains found in Tanzania, is required (Quibod et al., 2020; Vera Cruz et al., 2000).

We hypothesized that editing all known EBEs for both African and Asian SWEET-targeting TALes could be used to engineer full and robust resistance in an emerging elite variety for Africa. Since this strategy requires simultaneous editing of six EBEs, optimally by small deletions inside the EBE to enhance robustness, we developed a hybrid CRISPR-Cas9/Cpf1 system that targets all six EBEs in Komboka. Both CRISPR-Cas9 and CRISPR-Cpf1 systems have been widely used in genome editing. SpCas9 is the first Cas version to offer high editing efficiency across a wide range of plant species. LbCpf1 or LbCas12 is another endonuclease which is smaller than SpCas9 and requires shorter CRISPR RNA (crRNA)(Liu et al., 2017). SpCas9 and LbCpf1 both generate double-stranded breaks, but Cas9 introduces blunted cuts 3 base pairs upstream of the PAM, while Cpf1 introduces 5-bp staggered cuts downstream of the PAM (Zetsche et al., 2015). Hence, Cpf1 often produces larger mutations compared to Cas9, which mainly introduces single base pair insertions or deletions (indels). Depending on the purpose of genome editing, large mutations or small mutations are preferred. To generate knock-out mutants, single base pair indels in the first exon that result in frameshifts or premature stop codons are sufficient. To prevent the binding of TALes to their respective EBEs on *SWEET* promoters, single base pair changes are not always effective due to the degenerate code and the ability to loop out central repeats of the TALes (Oliva et al., 2019). In addition, *Xoo* evolves rapidly and, therefore, single base pair changes could be overcome quickly, as demonstrated for an *xa25-like* resistance gene (Xu et al., 2019). Hence, to prevent rapid break of the resistance by simple modifications of the TAL effectors, a larger mutation would be preferred. Since it is still unknown how the native *SWEET* expression is affected by promoter mutations, the number of base pair indels needs to be evaluated carefully. In addition, prevention of TALe binding to EBEs depends not only on the number of deleted or inserted base pairs but also on the position of the introduced mutations within the EBE. Especially, the RVDs at the 3’-end of TALes, such as N* and NS, have less specificity in nucleotide binding, therefore, mismatches at the 3’-end of an EBE can be tolerated by the RVDs (Richter et al., 2014). Here, the 1-bp deletion at the 5’-end of the TalC binding site is sufficient for resistance, while a 4-bp deletion at the 3’-end of the AvrXa7 binding site did not prevent binding of AvrXa7 to its EBE, indicating that mutations at the 5’-end of the TALe binding site are more effective than mutations at the 3’-end in preventing the binding of TALes to EBEs.

Cas9 cleaves target DNA adjacent to a G-rich PAM, while Cpf1 requires a T-rich PAM sequence (Zetsche et al., 2015; Zhang et al., 2014). Therefore, combining Cas9 and Cpf1 into a single system enables simultaneous editing in both G- and T-rich regions. Here, EBE regions used for gRNA design were comparatively short, just 23-29 bp, hence, there were a limited number of suitable gRNA design options (considering also potential unwanted off-target effects). The BB-resistant IR64 and Ciherang-Sub1 lines developed previously for Asia were generated using four gRNAs for *SWEET11a, 13* and *14* promoters, but did not target the TalF EBE in *SWEET14* which is targeted by African strains specifically (Eom et al., 2019; Oliva et al., 2019). The EBE in the respective promoters for TalF lacked a suitable SpCas9 NGG-PAM recognition sequence. By combining the two enzymes Cpf1 and Cas9 into a hybrid CRISPR-Cas9/Cpf1 system, it became possible to design four gRNAs (three gRNAs for Cpf1, including one for TalF; and one for to target TalC, which lacked a suitable PAM sequence for Cpf1) that target all six known EBEs relevant for BB susceptibility. The combination of CRISPR-Cas9 and Cpf1 gRNAs in a single system increases the flexibility of choosing suitable gRNAs depending on the available PAM recognition sequences and off-target frequency. An efficient transformation protocol previously established for Komboka was used here to edit all known EBEs in three *SWEET* promoters (Luu et al., 2020). For four target sites (cXo1, cXo2c, gTalC, cTalF), we generated 15, 27, 16 and 21 variants, respectively using four gRNAs, demonstrating the successful application of the hybrid Cas9/Cpf1 in multiplex genome editing of the rice variety Komboka. Each variant represents a distinct *R* gene. Edited Komboka lines were shown to be resistant against a set of representative Asian and African *Xoo* strains, which harbor all known *SWEET*-inducing TALes.

Crosses and analyses for the reliable elimination of transgenes from the edited Komboka lines is ongoing. Ideally, the deployment of edited lines should be region-specific, based on the variation of the TALes present in the *Xoo* populations. In parallel, monitoring the evolution of *Xoo* in Africa is extremely necessary for early detection of Asian strains or hybrid African-Asian strains. In addition, the diagnostic SWEET^R^ Kit 2.0, upgraded with the newly identified *SWEET11b* susceptibility gene and containing all six translational SWEET-reporter and knockout lines, can effectively be used to detect the targeting EBEs, while TALome analysis should be performed to detect the presence of *SWEET*inducing TALes in the current African *Xoo* population (Eom et al., 2019; Wu et al., 2022). Kenya evaluates applications for the import of transgene-free lines on a case-by-case basis, thus it will be necessary to prepare suitable documentation for import. In parallel, it is planned to use the hybrid Cas9/Cpf1 strategy to expand the resistant germplasm to other rice varieties grown in African countries to be able to meet consumer-farmer preferences. This project aims to ultimately provide the material to small-scale producers. It will also be necessary to continue monitoring disease and *Xoo* populations. Governmental action is recommended to reduce the risk of introduction of Asian pathogens to Africa and vice versa. In parallel, improved management practices and training may also help farmers to reduce disease incidence (Mew et al., 2018).

## Methods

### Rice seeds and *Xoo* strains

Seeds of the rice variety Komboka (IR05N221, L17WS.06#24) were provided by the International Rice Research Institute (IRRI, The Philippines) under a Standard Material Transfer Agreement under the Multilateral System (SMTA-MLS). *Xoo* strains were obtained from the *Xoo* strain collection of at the French National Research Institute for Sustainable Development (IRD, Montpellier, France).

### Rice cultivation

Rice seed germination and plant cultivation at HHU were done as described (Luu et al., 2020). Briefly, rice seeds were sterilized and germinated onto ½ Murashige Skoog media, supplemented with 1% sucrose. The seedlings were grown for 10 days in magenta boxes before transferring to soil. The plants were grown in greenhouses maintained at 30 °C day/ 25 °C night at a relative humidity (RH) of 50-70% and with supplemental LED light (Valoya, BX100 NS1) using a 8-hr/16-hr day/night photoperiod at 400 μmol/m^-2^s^-1^. The plants were fertilized weekly from the 2^nd^ week and biweekly from the 6^th^ week after germination. At IRD, plants were sown in soil complemented with 3 g of fertilizer (N: 19%; P: 5%; K: 8%) per liter and grown in greenhouses (12-h light at 200 μmol/m^-2^s^-1^, 28 °C, 80% RH; and12-h darkness at 25 °C and 60% RH).

### Disease resistance scoring by leaf-clipping inoculation

Bacteria were grown on PSA media (Peptone Sucrose Agar, 1% peptone, 1% sucrose, 0.1% glutamic acid, 1.6% agar) for four days at 28 °C. Single colonies were picked and patched onto PSA and then grown for 24 h before being washed with and diluted in sterile water at OD_600_ 0.2 (at IRD) and 0.5 (at HHU). The youngest, fully extended leaves of 4-to 6-week-old rice plants were clipped 2-3 cm from the leaf tip with scissors that had been dipped in the inoculum or sterile water. Lesion lengths were measured 14 days after inoculation. For the race typing assay, lesion length measurements < 5 cm were scored as resistant (R), 5–10 cm as moderately resistant (MR), 10–15 cm as moderately susceptible (MS), and >15 cm as susceptible (S).

### Sampling of symptomatic leaf material and isolation of *Xoo* strains

Small-scale rice fields cultivated under irrigated conditions in the villages of Dakawa and Lukenge (in the Morogoro region of Tanzania) were surveyed for BB between 2019 and 2022. In each field, diseased leaves of several individual plants were sampled and processed for bacterial isolation as reported previously (Tekete et al., 2020). Colony-multiplex PCR was used to validate that isolates were *Xanthomonas oryzae* pv. *oryzae* (except for 2022 data, for which analyses have been initiated).

### Tanzanian strain genome sequencing and analysis

DNA was extracted from pure bacterial cultures grown on rich media (1% peptone, 0.1% glutamic acid) with QIAGEN Genomic-tip 100/G. Multiplex libraries were prepared with the rapid library preparation kit (SQK-RBK110-96, Oxford Nanopore Technologies, ONT) and sequenced with a MinION Mk1C device on R10.3 (FLO-MIN111) flow cells. ONT electric signals were base-called with a high accuracy model (dna_r10.3_450bps_hac.cfg) and demultiplexed with the ONT Guppy base calling software (v6.0.1+652ffd179). Illumina library construction and sequencing was performed by FASTERIS (Plan-les-Ouates, Switzerland) on an Illumina NextSeq sequencer with 350-bp inserts and 150-bp paired-end sequences. Long ONT read assembly was performed with the CulebrONT pipeline (v2.0.1)(Orjuela et al., 2022) and included FLYE (2.9-b1768) for primary assembly followed by RACON (v1.4.20) and MEDAKA (1.4.1) for assembly polishing. The ONT-only assemblies were subsequently polished with polypolish (v0.5.0)(Wick and Holt, 2022). Core genome SNP genotyping and tree reconstruction were conducted as previously described (Doucouré et al., 2018), except that the raxml command included “-N 500 −m GTRGAMMAIX”. This substitution model was adopted based on the output of the ‘modelTest’ function of the R package phangorn for model selection (Schliep, 2011). The X11-5A assembly (GB Acc. GCF_000212755.2) was used as an out-group, and was not displayed in Figure 2. The genomes included in the species-wide *Xanthomonas oryzae* phylogenetic tree are listed together with NCBI accessions and genome metadata in Supplementary Data 1 and Supplementary Table 3. Predictions for *TALe* genes from the genomes of the Tanzanian strains used AnnoTALE with default parameters (Grau et al., 2016). Predictions for high-scoring EBEs of the Tanzanian TALes in the Nipponbare *SWEET11a* promoter used Talvez with default parameters (Pérez-Quintero et al., 2013). Talvez target predictions on the Nipponbare SWEET11a promoter included putative TALes from Tanzanian strains (CIX4462, CIX4506, CIX4509, T19, Dak16, and Xoo3-1), Asian strains (PXO99A, PXO86, PXO83, PXO71, PXO61) and an African strain (MAI1), which were included as references. Only predictions with a score above 9.05 are displayed in Figure 2.

### qRT-PCR analyses of *SWEET11a* mRNA levels

Leaves of 3-week-old rice plants were infiltrated with a bacterial suspension at an OD_600_ of 0.5 or water using a needleless syringe. One leaf per plant was infiltrated, and three plants per treatment were used. Samples were collected at 24 h (Kitaake) or 48 h (wildtype Komboka and edited lines) after inoculation. Total RNA was extracted using TRI reagent (Euromedex). Following extraction, DNase I treatment was conducted using Turbo DNA-free kit (Thermo Fisher Scientific). Subsequent synthesis of complementary DNA was carried out using SuperScriptIII (Thermo Fisher Scientific) and oligo-dT primers. Quantitative PCR reactions were performed with SYBR Mesa Blue qPCR Mastermix (Eurogentec). Three technical replicates were prepared for each sample. Expression values were normalized by subtracting the values obtained for reference gene *EF-1α* (GenBank: GQ848072.1) to the studied replicates (2^-ΔCt^ method). Mean of the technical replicates for each sample was calculated. Three independent biological replicates were analyzed. Primer sequences are listed in Supplementary Table 7.

### Analysis of EBEs in *SWEET* promoters of *O. sativa* cv. Komboka

Genomic DNA was extracted from Komboka leaves using the CTAB method (Li et al., 2013). The gDNA fragments corresponding to promoters and first exons of *SWEET11a, 13, 14* were PCR-amplified using specific primer pairs. Amplicons were gel-purified and cloned into pJET1.2 (Thermo Fischer Scientific). Competent *E.coli* TOP10 competent cells were used for transformation. Plasmids were extracted from three individual colonies and were sent for Sanger sequencing. Sanger sequencing reads representing promoter regions of *SWEET11a, 13*, and *14* from Komboka were aligned with the respective promoter regions of Kitaake (Kitaake_OsativaKitaake_499.genome; Phytozome v.13) (Supplementary Data 2).

### Analyses of *Xa1* and *Xa4* sequences from *O. sativa* cv. Komboka

Full-length *Xa1* and four fragments of *Xa4* were PCR-amplified from *O. sativa* cv. Komboka gDNA using TAKARA PrimeSTART GXL polymerase and primers. Amplicons corresponding to full length *Xa1* were subjected to Sanger sequencing. *Xa4* fragments were subcloned into pUC57Gent vector (Addgene #54338)(Binder et al., 2014) and sequenced using Sanger sequencing. Individual sequence reads were mapped to their references and assembled via Geneious Prime^®^ 2022.1.1. Neighbor-joining phylogenetic analyses were made using CLUSTAL Omega (Geneious Prime). Genomic sequences of non-Komboka derived *Xa1* and *Xa4* had previously been published (Hu et al., 2017; Ji et al., 2020). Sequences of *Xa1* and *Xa4* genes used for phylogenetic analysis are provided in Supplementary Data 3.

### Hybrid CRISPR/Cas9 and Cpf1 vector construction

A new hybrid CRISPR-Cas9/Cpf1 system was developed to edit a maximum of 6 different targets (Figure 7-supplement 2). The hybrid CRISPR-Cas9/Cpf1 combines a duplex Cas9 gRNA combined with a multiplex Cpf1 cRNA. To develop this hybrid CRISPR-Cas9/Cpf1 system, we subcloned a human-codon-optimized *LbCpf1*-coding sequence (www.addgene.org/69988/sequences) (Zetsche et al., 2015) and the rice Ubiquitin 1 (*OsUbi*) (LOC_Os02g06640) to create a gateway system pDEST:Cpf1. A PCR fragment of a Cas9 expression cassette, which contains a rice-codon-optimized *SpCas9*-coding sequence driven by the maize Ubiquitin 1 (*ZmUbi*) promoter (Char et al., 2017; Zhou et al., 2014), was amplified and inserted into pDEST:Cpf1 by Gibson Assembly (New England BioLabs). Finally, we constructed the hybrid destination vector pDEST:Cpf1&Cas9. A Ribozyme-gRNA-Ribozyme (RGR) system was used to generate multiple cRNAs with different target sequences by flanking the cRNAs with a Hammerhead (HH) type ribozyme and a hepatitis delta virus (HDV) ribozyme (Gao and Zhao, 2014). Guide RNA sequences are provided in Supplementary Table 8. The promoter of the small nuclear RNA gene from rice *OsU6.1* and *OsU3*, wheat *TaU3* and maize *ZmU3* were used to drive expression of RGG-cRNAs units. Four gBlock fragments synthesized by IDT (Integrated DNA Technologies, Inc., Iowa, USA) were inserted into the vector pTLN using *XbaI* and *XhoI* to produce four intermediate pUNIT vectors: pTL-OsU6.1_CpfRNA1, pTL-OsU3_CpfRNA2, pTL-TaU3_CpfRNA3, pTL-ZmU3_CpfRNA4. A double-stranded DNA oligonucleotide for each site was produced by annealing two complementary oligonucleotides. The DNA sequence of positive clones was confirmed by Sanger sequencing. All four pUNIT vectors were transferred into another donor vector named pENTR4-ccdB using Golden Gate cloning to generate pENTR:cRNAs containing Cpf1 gRNAs cXo1, cXo2c, cTalF and cXo2d. Simultaneously, a gTalC double-stranded oligonucleotide was inserted into the *BtgZI-digested* pENTR-gRNAs (Char et al., 2017). The gRNA expression cassette was PCR-amplified with oligos U6P-F3 and -R3 inserted into pENTR:gRNAs at *XbaI* site through Gibson cloning to generate the hybrid donor plasmid pENTR:gRNA-cRNAs. The gRNA-cRNA cassette was mobilized to the hybrid destination vector pDEST:Cpf1&Cas9 using Gateway LR Clonase (Thermo Fisher Scientific) to produce a pCam1300-CRISPR plasmid named pCpf1&Cas9:gRNAs-cRNAs (pMUGW5). DNA sequencing of the plasmid pMUGW5 detected the insertion of a *E. coli* IS element in the vector backbone. The insertion was already present in the original pCambia3000 in BY stock collection and apparently did not affect transformation or editing.

### Rice transformation

*Agrobacterium-mediated* transformation of Komboka using immature embryos was performed as described (Luu et al., 2020). Briefly, the *Agrobacterium tumefaciens* strain LBA4404 was transformed with pMUGW5 via electroporation. Immature rice seeds at the late milky stage were harvested for immature embryo isolation. Immature embryos were inoculated with 5 μl *Agrobacterium* suspension (OD_600_ 0.3) and incubated in the dark for one week. The emerged shoots were removed, and immature embryos were incubated for 5 days at 28 °C under continuous light. To select positive transformants, four rounds of hygromycin selection were applied, with each round consisting of 10 days. Resistant calli were moved onto a pre-regeneration medium and incubated for 10 days. Greening calli were transferred to a regeneration medium to develop small rice plantlets. Once plantlets reached 15-cm in height, they were transferred to soil and placed in a greenhouse. After 5 months, T1 seeds were harvested.

### Screening of EBE edited lines

To test whether generated T0 plants contained T-DNA insertion, the presence of *SpCas9, LbCpf1* and hygromycin resistance (*Hpt*) genes were checked. Leaf fragments (3 to 4 cm) were harvested for DNA extraction using a modified CTAB method for high-throughput DNA extraction. PCR was performed using GoTaq DNA Polymerase (Promega) with a melting temperature of 55 °C for *SpCas9, LbCpf1* and *Hpt*. For genotyping of EBE mutations, the four regions containing the six EBEs within the *SWEET11a*, *13* and *14* promoters were amplified with specific primers using Phusion HF polymerase (Thermo Fisher Scientific), and PCR amplicons were sequenced by Sanger sequencing (Microsynth seqlab). Chromatograms were analyzed manually using Benchling (www.benchling.com) to detect the mutations. Two rounds of transformation were performed. In Round 1, seven T0 plants were obtained from a single embryo, and all T0 plants contained biallelic mutations at all target sites (Supplementary Table 5, Supplementary Table 9, Supplementary Table 10). In the transformation Round 2, 18 T0 plants were T-DNA positive, from 18 independent immature embryos. Nine T0 plants were characterized further and found to contain biallelic mutations at all targeted sites (Supplementary Table 10 Supplementary Table 12,). We screened 52 T1 plants from three independent events (#12, 14 and 16) and obtained 7 lines with homozygous mutations in all 6 EBEs (Supplementary Table 5, Supplementary Table 6, Supplementary Table 9).

### Statistical analyses

Statistical analyses and graphical representations were prepared using R 4.0.5, on RStudio for Windows. Statistics were calculated using rstatix (https://cran.r-project.org/web/packages/rstatix). To determine if there was an effect of the treatment or the genotype on gene expression or lesion length, the medians of the different subsamples were compared using rstatix Kruskal-Wallis test (*p*-values < 0.05). Then, mean groups were attributed to the different genotypes or treatments independently using rstatix Dunett’s test (*p*-values < 0.05). Letters were attributed to each mean group. Graphical representations were realized using package ggplot2 (https://ggplot2.tidyverse.org). Data were drawn as boxplots. The boxplot is delimited by the first and the third quartile of the distribution of the studied variable. The line inside the boxplot represents the median of the variable. Finally, the two lines that start from the boxplot join the minimum and maximum theoretical values. Outliers (<7% of total values) were represented as black dots. The total number (n) of replicates for each strain/genotype condition is displayed next to each boxplot. Each observation is represented by a dot, with a color code by strain or genotype. The letters above each boxplot represent the mean groups calculated using Dunnett’s test.

## Acknowledgments

We thank Paula Emmerich Maldonado (HHU) and Britta Killing (HHU) for excellent technical assistance. We thank Dr. Marietta Wolter for advice on using the QX200 Droplet Digital PCR (ddPCR) System (Biorad) at the Institute for Neuropathology, University Hospital Düsseldorf. The authors acknowledge the ISO 9001-certified IRD i-Trop HPC (South Green Platform) at IRD Montpellier for providing high-performance computing resources. WBF and YA acknowledge support by the Alexander von Humboldt Foundation. This work was made possible by funding from the Bill and Melinda Gates Foundation to Heinrich Heine University, Düsseldorf, with subawards to Iowa State University/University of Missouri, University of Florida, Institut de Recherche pour le Développement, and the International Rice Research Institute.

## Data availability

All data supporting the results are available in the main text or supplementary materials. All data that support the findings of this study are available from the corresponding author upon request. Sequencing data for strains from this study have been deposited in the NCBI Sequence Read Archive (SRA) database under the accession codes will be provided at the time of publication. Source data will be deposited before publication. Materials will be made available under MTA.

## Contributions

V.S.L., M.S., Y.A., K.S., M.B., and E.P.I.L.: *R*-genes in Komboka, editing, genotyping; C.S., G.B. and F.A.: Race-typing, characterization of the resistance of genome-edited lines; F.A., C.S., A.D. and S.C.: strain isolation, whole genome sequencing and analyses; C.J., S.N.C., and B.Y.: hybrid Cas9/Cpf1 system; A.L.B., D.L., M.M., and R.M.: survey in Tanzania, disease monitoring and sampling; R.O., B.Y., B.S., and W.B.F.: conceived and designed experiments; V.S.L., M.S., Y.A., E.P.I.L., C.S., S.C, R.S., B.Y., B.S., and W.B.F. analyzed the data; R.O., M.B., B.Y., B.S. and W.B.F. wrote the manuscript.

## Ethics declarations

### Competing interests

The <u>Healthy Crops</u> team has filed for patents, but is charitable and non-profit and aims at helping small scale rice farmers in Asia and Africa by reducing yield losses caused by pathogens. The authors declare no competing interests.

---

**Figure 1-supplement 1.**
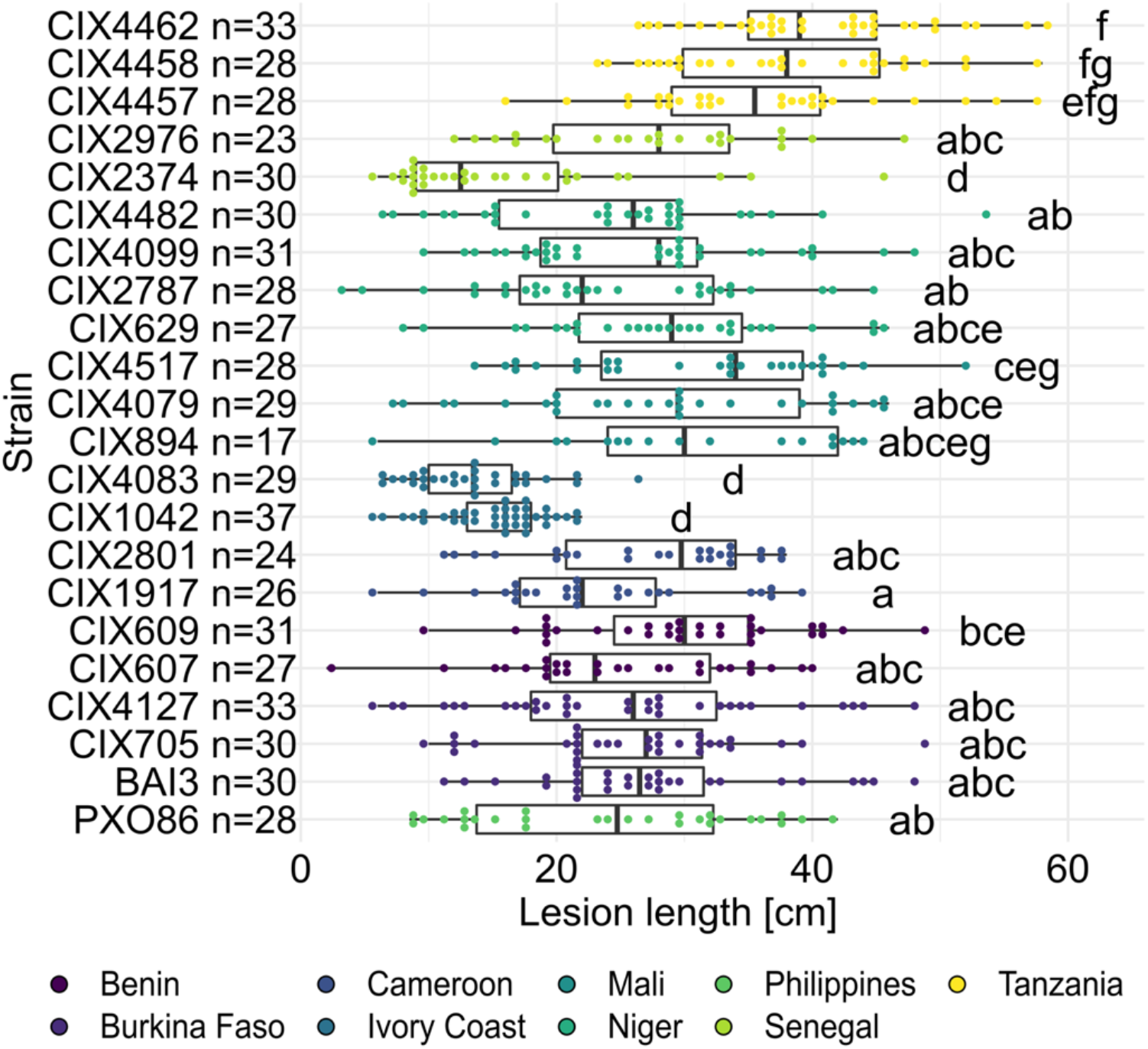
Virulence of the African *Xoo* diversity panel on the susceptible rice line Azucena. Leaf-clip inoculation of wildtype Azucena with a panel of 21 *Xoo* strains originating from 8 African countries along with Asian reference strain PXO86. Lesion lengths were measured 14 days after inoculation. Data from three independent experiments are represented.

**Figure 1-supplement 2.**
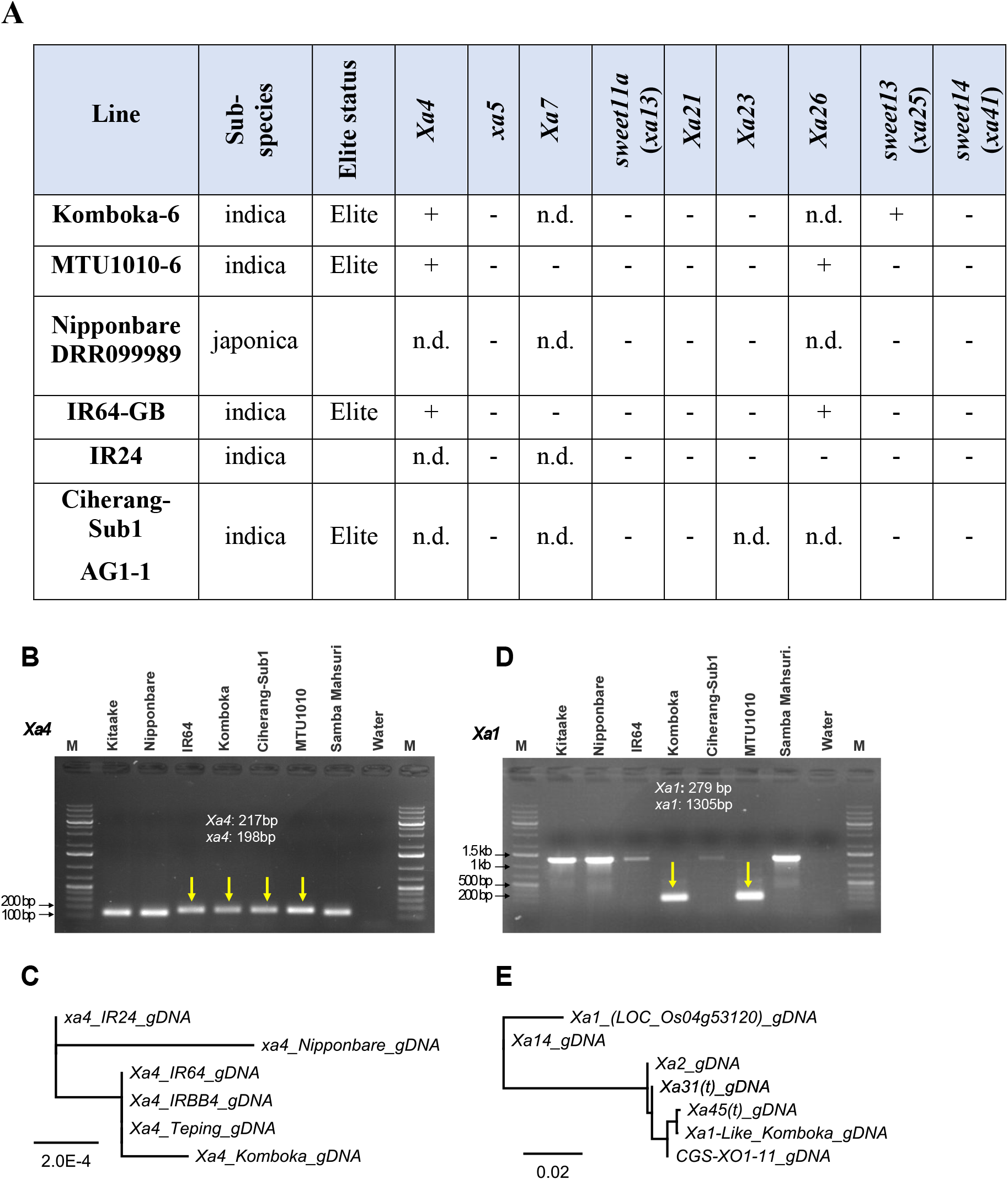
Presence of bacterial blight *R* genes in select elite rice varieties. **A.**QTL profiling of Komboka and other selected varieties from IRRI QTL database (https://rbi.irri.org/resources-and-tools/qtl-profiles). +: presence, −: absence, n.d., not determined. **B.-D. Presence of *Xa1* and *Xa4 R* genes in Komboka using their associated markers and sequencing. B.**Genotyping of *O. sativa* cultivars for an *Xa4*-associated marker gene. **C.**Neighbor-joining phylogenetic analysis of *Xa4* from *O. sativa* cultivars. *Xa4* from Komboka shares 99.98% identity with *Xa4* from IR64, Teping, and IRBB4 (Hu et al., 2017). **D.**Genotyping of *O. sativa* cultivars for presence of functional *Xa1-like* genes with gene-specific markers. **E.**Neighbor-joining phylogenetic analysis of *Xa1* from *O. sativa* cultivars. Komboka *Xa1* is 99.88% homologous to *Xa45(t*) (Ji et al., 2020).

**Figure 4-supplement 1.**
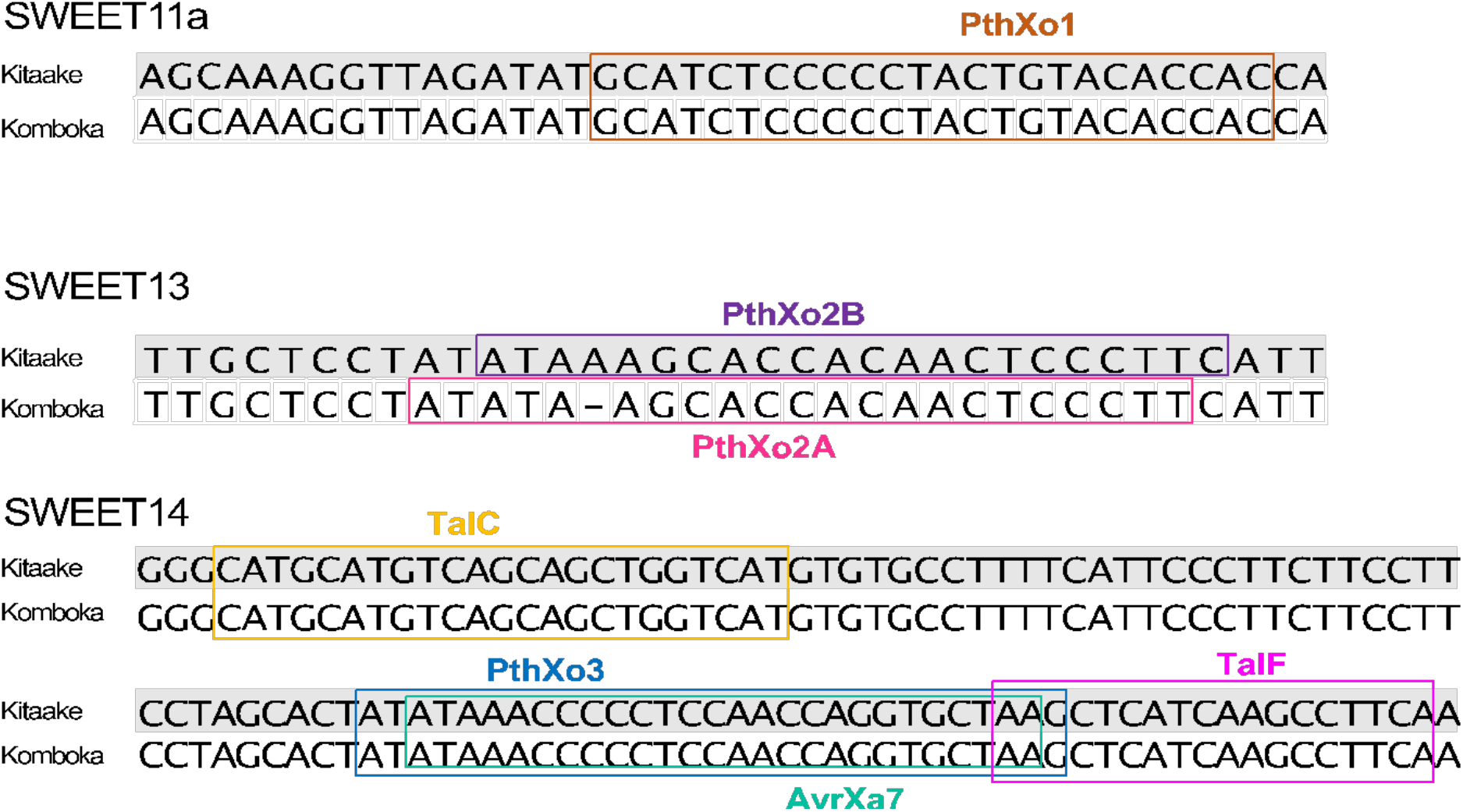
Comparison of EBE sequences for the *SWEET11a, 13* and *14* promoters between Komboka and Kitaake. Kitaake and Komboka have the same EBE sequences for PthXo1, TalC, TalF, AvrXa7, PthXo3. Komboka contains the EBE for PthXo2A, while Kitaake contains the EBE for PthXo2B in the *SWEET13* promoter.

**Figure 4-supplement 2.**
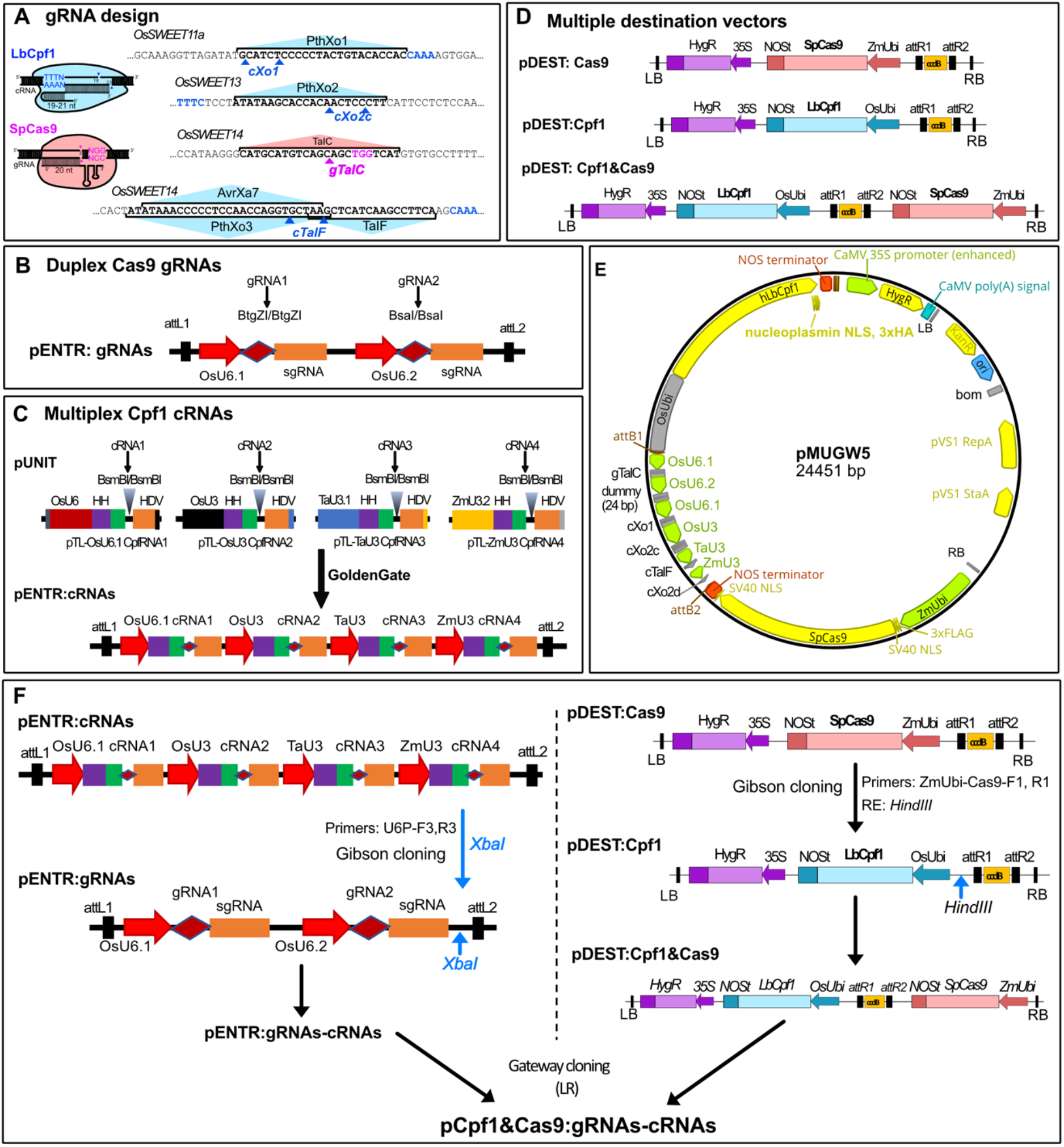
Hybrid CRISPR-Cas9/Cpf1 system. **A**. Guide RNA design. Four guide RNAs were designed to target six known EBEs within *OsSWEET11a, 13* and *14* promoters. EBE sequences are bolded, and the respective TALEs are indicated. Arrowheads indicate guide nuclease cleavages sites. Cas9 guide RNA is colored in pink (gTalC), and Cpf1 guide RNAs are colored in blue (cXo1, cXo2c, cXo2d, cTalF) (cX02d is for another *SWEET13* allele). **B**. Duplex CRISPR-Cas9 guide RNA system. Single Cas9 gRNA were cloned into the pENTR:gRNA. **C**. Multiplex CRISPR-Cpf1 guide RNA system. Single Cpf1 cRNA were shuttled into the unit vectors and assembled into pENTR:cRNAs via GoldenGate assembling. **D**. Three types of destination vectors: pDEST:Cas9 is the acceptance vector of pENTR:gRNAs, pDEST:Cpf1 is the acceptance vector of pENTR:cRNAs and hybrid pDEST:Cpf1&Cas9 is the acceptance vector for both Cas9 and Cpf1 guide RNAs. **E.**pMUGW5 map that contains 4 Cpf1 gRNAs and 1 Cas9 gRNA targeting six EBEs on *OsSWEET11a, 13* and *14*. **F.**Cloning scheme. The PCR fragment containing a rice-codon-optimized *SpCas9*-coding sequence driven by the maize Ubiquitin 1 (*ZmUbi*) promoter from pDEST:Cas9 was amplified and inserted into pDEST:Cpf1 to create a hybrid destination vector pDEST:Cpf1&Cas9. Four gBlock fragments were inserted into the vector pTLN using *Xba*I and *Xho*I to generate four intermediate pUNIT vectors: pTL-OsU6.1-CpfRNA1, pTL-OsU3-CpfRNA2, pTL-TaU3-CpfRNA3, pTL-ZmU3-CpfRNA4. All four pUNITs, after cloning of respective guide RNAs, were transferred into the donor vector pENTR4-ccdB to generate pENTR:cRNAs containing the Cpf1 gRNAs cXo1, cXo2c, cTalF and cXo2d. Simultaneously, gTalC double-stranded oligonucleotides was inserted into the *BtgZ*I-digested pENTR-gRNA. The cRNA expression cassette was PCR-amplified and inserted into pENTR:gRNAs at the *Xba*I site through Gibson cloning to generate the hybrid donor plasmid pENTR:gRNA-cRNAs. The gRNA and cRNAs cassettes were mobilized to the hybrid destination vector pDEST:Cpf1&Cas9 by Gateway to create a pCam1300-CRISPR plasmid named pCpf1&Cas9:gRNAs-cRNAs (pMUGW5).

**Figure 4-supplement 3.**
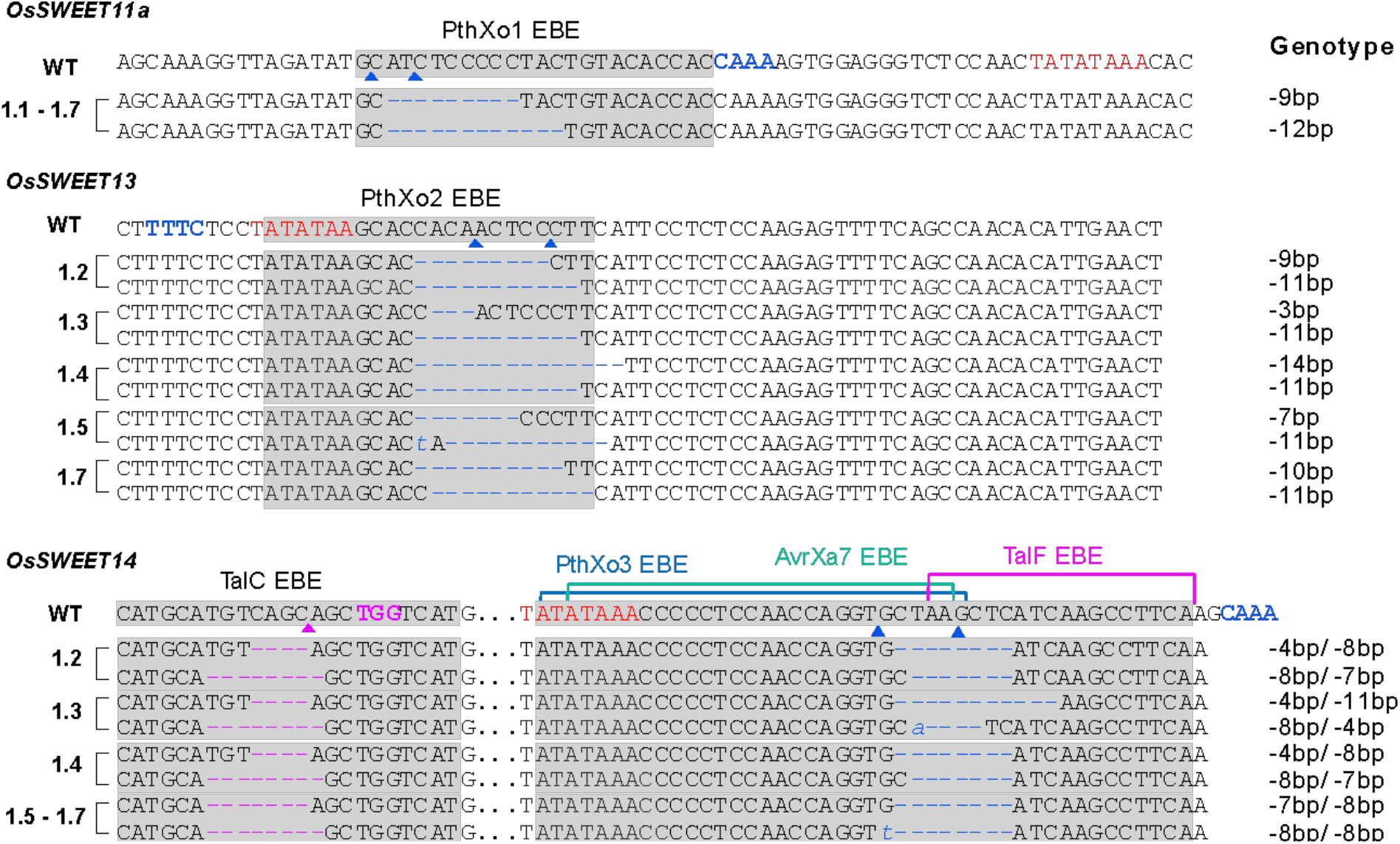
Genotypes of T0 lines with biallelic mutations at six targeted EBEs. TATAA box is labeled in red. LbCpf1 and SpCas9 PAM sequences are labeled in blue and purple, respectively. EBE sequences are indicated (e.g., PthXo1).

**Figure 6-supplement 1.**
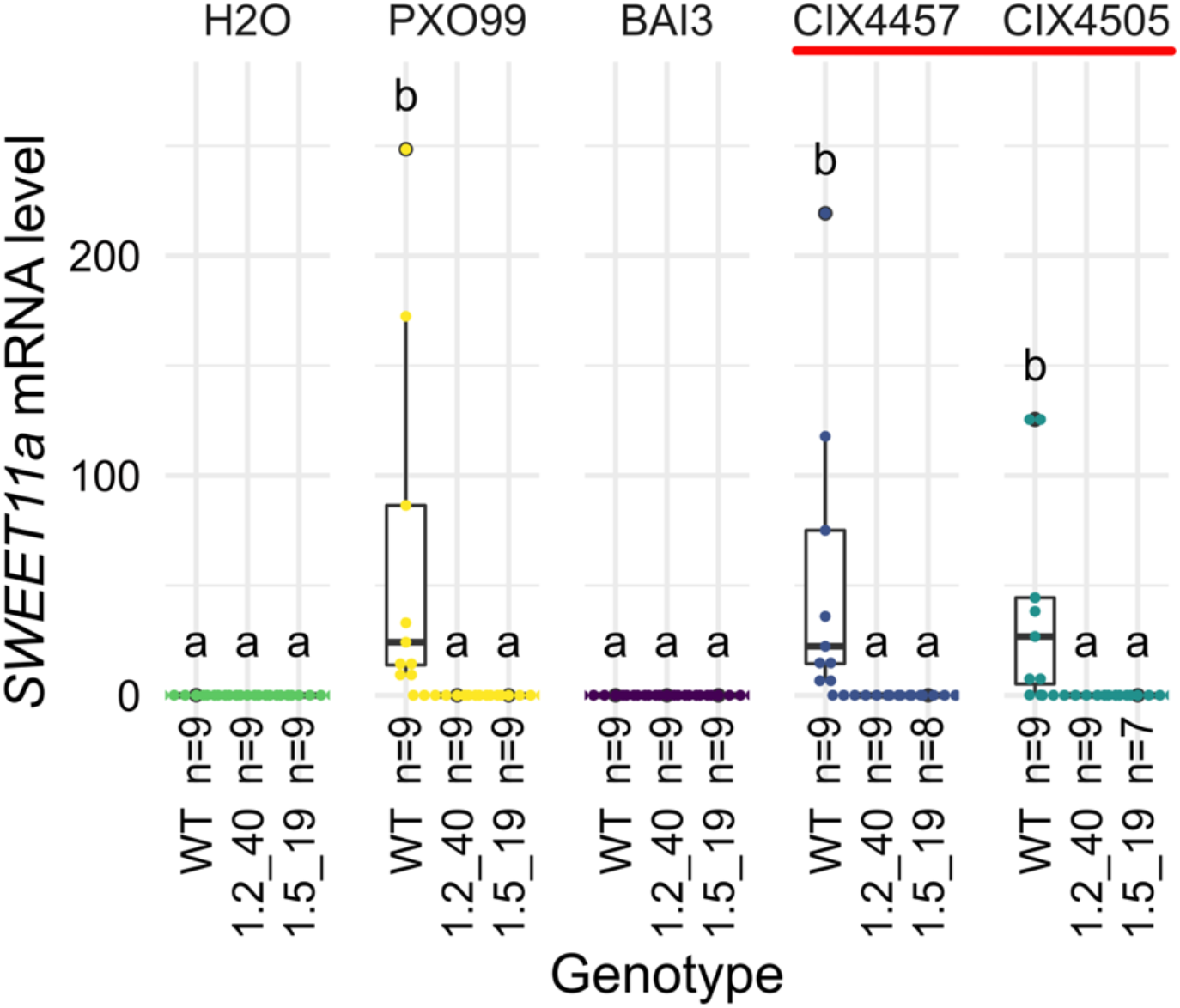
qRT-PCR analysis of *SWEET11a* mRNA accumulation of edited Komboka lines after infection with Tanzanian *Xoo* strains. Relative mRNA levels (2^-ΔCt^) of *SWEET11a* in Komboka wild type and the multi-EBE-edited lines 1.2_40 and 1.5_19 upon infection by PXO99 (PthXo1), BAI3 (TalC) and Tanzanian *PthXo1B* dependent *Xoo* strains (red line). Samples were collected 48h post infiltration. Three independent experiments are represented. Three technical replicates were performed for each sample. All Ct values were normalized by rice *EF1α* elongation factor (△Ct).

## Notes

### Competing Interest Statement

The authors have declared no competing interest.

## References

Afolabi O, Amoussa R, Bilé M, Oludare A, Gbogbo V, Poulin L, Koebnik R, Szurek B, Silué D. 2016. First report of bacterial leaf blight of rice caused by *Xanthomonas oryzae pv. oryzae* in Benin. Plant Dis 515.

Amos O. 2013. Two genotypes of *Xanthomonas oryzae pv. oryzae* virulence identified in West Africa. Int J Genet Mol Biol 5:28–36. doi:10.5897/IJGMB2013.0070

Arra Y, Sundaram RM, Ladhalakshmi D, Hajira SK, Prakasam V, Prasad MS, Sheshu Madhav M, Ravindra Babu V, Laha GS. 2017. Virulence profiling of *Xanthomonas oryzae pv. oryzae* isolates, causing bacterial blight of rice in India. Eur J Plant Pathol 149:171–191. doi:10.1007/s10658-017-1176-y

Arra Y, Sundaram RM, Singh K, Ladhalakshmi D, Subba Rao LV, Madhav MS, Badri J, Prasad MS, Laha GS. 2018. Incorporation of the novel bacterial blight resistance gene *Xa38* into the genetic background of elite rice variety improved Samba Mahsuri. PloS One 13:e0198260. doi: 10.1371/journal.pone.0198260

Binder A, Lambert J, Morbitzer R, Popp C, Ott T, Lahaye T, Parniske M. 2014. A modular plasmid assembly kit for multigene expression, gene silencing and silencing rescue in plants. PLOS ONE 9:e88218. doi:10.1371/journal.pone.0088218

Buchholzer M, Frommer WB. 2022. An increasing number of countries regulate genome editing in crops. New Phytol online. doi:10.1111/nph.18333

Buddenhagen IW, Vuong HH, Ba DD. 1979. Bacterial blight found in Africa. Int Rice Res Newsl 4:11.

Char SN, Neelakandan AK, Nahampun H, Frame B, Main M, Spalding MH, Becraft PW, Meyers BC, Walbot V, Wang K, Yang B. 2017. An Agrobacterium-delivered CRISPR/Cas9 system for high-frequency targeted mutagenesis in maize. Plant Biotechnol J 15:257–268. doi:10.1111/pbi.12611

Chen LQ, Qu X-Q, Hou B-H, Sosso D, Osorio S, Fernie AR, Frommer WB. 2012. Sucrose efflux mediated by SWEET proteins as a key step for phloem transport. Science 335:207–211. doi:10.1126/science.1213351

Doucouré H, Pérez-Quintero AL, Reshetnyak G, Tekete C, Auguy F, Thomas E, Koebnik R, Szurek B, Koita O, Verdier V, Cunnac S. 2018. Functional and genome sequence-driven characterization of TAL effector Gene repertoires reveals novel variants with altered specificities in closely related Malian *Xanthomonas oryzae pv. oryzae* strains. Front Microbiol 9:1657. doi:10.3389/fmicb.2018.01657

Duku C, Sparks AH, Zwart SJ. 2016. Spatial modelling of rice yield losses in Tanzania due to bacterial leaf blight and leaf blast in a changing climate. Clim Change 135:569–583. doi:10.1007/s10584-015-1580-2

Eom J-S, Luo D, Atienza-Grande G, Yang J, Ji C, Luu VT, Huguet-Tapia JC, Liu B, Nguyen H, Schmidt SM, Szurek B, Vera-Cruz CM, White FF, Oliva R, Yang B, Frommer WB. 2019. Diagnostic kit for rice blight resistance. Nat Biotechnol 37:1372–1379.

Gao Y, Zhao Y. 2014. Self-processing of ribozyme-flanked RNAs into guide RNAs in vitro and in vivo for CRISPR-mediated genome editing. J Integr Plant Biol 56:343–349. doi:10.1111/jipb.12152

Gonzalez C, Szurek B, Manceau C, Mathieu T, Séré Y, Verdier V. 2007. Molecular and pathotypic characterization of new *Xanthomonas oryzae* strains from West Africa. Mol Plant-Microbe Interact MPMI 20:534–546. doi: 10.1094/MPMI-20-5-0534

Grau J, Reschke M, Erkes A, Streubel J, Morgan RD, Wilson GG, Koebnik R, Boch J. 2016. AnnoTALE: bioinformatics tools for identification, annotation, and nomenclature of TALEs from Xanthomonas genomic sequences. Sci Rep 6:21077. doi:10.1038/srep21077

Hu K, Cao J, Zhang J, Xia F, Ke Y, Zhang H, Xie W, Liu H, Cui Y, Cao Y, Sun X, Xiao J, Li X, Zhang Q, Wang S. 2017. Improvement of multiple agronomic traits by a disease resistance gene via cell wall reinforcement. Nat Plants 3:1–9. doi:10.1038/nplants.2017.9

Ji C, Ji Z, Liu B, Cheng H, Liu H, Liu S, Yang B, Chen G. 2020. Xa1 allelic R genes activate rice blight resistance suppressed by interfering TAL effectors. Plant Commun, Special Issue on Plant-Pathogen Interactions (Organizing Editors: Paul Birch, Savithramma Dinesh-Kumar, Hui-Shan Guo, Ping He, Xin Li, Frank Takken, Yuanchao Wang) 1:100087. doi:10.1016/j.xplc.2020.100087

Jiang N, Yan J, Liang Y, Shi Y, He Z, Wu Y, Zeng Q, Liu X, Peng J. 2020. Resistance genes and their interactions with bacterial blight/leaf streak pathogens (*Xanthomonas oryzae*) in rice (*Oryza sativa L.*)—an updated review. Rice 13:3. doi:10.1186/s12284-019-0358-y

Kitilu MJF, Nyomora AMS, Charles J. 2019. Growth and yield performance of selected upland and lowland rainfed rice varieties grown in farmers and researchers managed fields at Ifakara, Tanzania. Afr J Agric Res 14:197–208. doi:10.5897/AJAR2018.13611

Lang JM, Hamilton JP, Diaz MGQ, Van Sluys MA, Burgos MaRG, Vera Cruz CM, Buell CR, Tisserat NA, Leach JE. 2010. Genomics-based diagnostic marker development for *Xanthomonas oryzae pv. oryzae* and *X. oryzae pv. oryzicola*. Plant Dis 94:311–319. doi:10.1094/PDIS-94-3-0311

Lang JM, Pérez-Quintero AL, Koebnik R, DuCharme E, Sarra S, Doucoure H, Keita I, Ziegle J, Jacobs JM, Oliva R, Koita O, Szurek B, Verdier V, Leach JE. 2019. A pathovar of *Xanthomonas oryzae* infecting wild grasses provides insight into the evolution of pathogenicity in rice agroecosystems. Front Plant Sci 10. doi:10.3389/fpls.2019.00507

Li H, Li J, Cong XH, Duan YB, Li L, Wei PC, Lu XZ, Yang JB. 2013. A high-throughput, high-quality plant genomic DNA extraction protocol. Genet Mol Res GMR 12:4526–4539. doi:10.4238/2013.October.15.1

Liu Y, Han J, Chen Z, Wu H, Dong H, Nie G. 2017. Engineering cell signaling using tunable CRISPR–Cpf1-based transcription factors. Nat Commun 8:2095. doi:10.1038/s41467-017-02265-x

Longue RDS, Traore VSE, Zinga I, Asante MD, Bouda Z, Neya JB, Barro N, Traore O. 2018. Pathogenicity of rice yellow mottle virus and screening of rice accessions from the Central African Republic. Virol J 15:6. doi:10.1186/s12985-017-0912-4

Luu VT, Stiebner M, Maldonado PE, Valdés S, Marín D, Delgado G, Laluz V, Wu L-B, Chavarriaga P, Tohme J, Slamet-Loedin IH, Frommer WB. 2020. Efficient Agrobacterium-mediated transformation of the elite–indica rice variety Komboka. Bio-Protoc 10:e3739–e3739.

Makundi H. 2017. Diffusing Chinese rice technology in rural Tanzania: Lessons from the Dakawa agrotechnology demonstration center (No. 2017/12), SAIS-CARI Working Papers, SAIS-CARI Working Papers. Johns Hopkins University, School of Advanced International Studies (SAIS), China Africa Research Initiative (CARI).

Mew TW, Hibino H, Savary S, Vera Cruz CM, Opulencia R, Hettel GP. 2018. Rice diseases: Biology and selected management practices. Los Baños (Philippines): International Rice Research Institute.

Mutiga SK, Rotich F, Were VM, Kimani J, Mwongera DT, Mgonja E, Onaga G, Konaté K, Razanaboahirana C, Bigirimana J, Ndayiragije A, Gichuhi E, Telebacnco-Yanoria MJ, Otipa M, Wasilwa L, Ouedraogo I, Mitchell T, Wang G-L, Correll J, Talbot N. 2021. Integrated strategies for durable rice blast resistance in sub-Saharan Africa. Plant Dis 105:2749–2770. doi:10.1094/PDIS-03-21-0593-FE

Odongo PJ, Onaga G, Ricardo O, Natsuaki KT, Alicai T, Geuten K. 2021. Insights into natural genetic resistance to rice yellow mottle virus and implications on breeding for durable resistance. Front Plant Sci 12:671355. doi:10.3389/fpls.2021.671355

Oliva R, Ji C, Atienza-Grande G, Huguet-Tapia JC, Pérez-Quintéro A, Li T, Eom J-S, Li C, Nguyen H, Liu B, Cunnac S, Slamet-Loedin IH, Vera Cruz CM, Szurek B, Frommer WB, White FF, Yang B. 2019. Broad-spectrum resistance to bacterial blight in rice using genome-editing. Nat Biotechnol 37:1344–1350.

Orjuela J, Comte A, Ravel S, Charriat F, Vi T, Sabot F, Cunnac S. 2022. CulebrONT: a streamlined long reads multi-assembler pipeline for prokaryotic and eukaryotic genomes. bioRxiv 2021.07.19.452922. doi:10.1101/2021.07.19.452922

Pandey S, Byerlee DR, Dawe D, Dobermann A, Mohanty S, Rozelle S, Hardy B. 2010. Rice in the global economy: strategic research and policy issues for food security, IRRI Books. International Rice Research Institute (IRRI).

Pérez-Quintero AL, Rodriguez-R LM, Dereeper A, López C, Koebnik R, Szurek B, Cunnac S. 2013. An improved method for TAL effectors DNA-binding sites prediction reveals functional convergence in TAL repertoires of *Xanthomonas oryzae* strains. PLOS ONE 8:e68464. doi: 10.1371/journal.pone.0068464

Quibod IL, Atieza-Grande G, Oreiro EG, Palmos D, Nguyen MH, Coronejo ST, Aung EE, Nugroho C, Roman-Reyna V, Burgos MR, Capistrano P, Dossa SG, Onaga G, Saloma C, Cruz CV, Oliva R. 2020. The Green Revolution shaped the population structure of the rice pathogen Xanthomonas oryzae pv. oryzae. ISME J 14:492–505. doi:10.1038/s41396-019-0545-2

Quibod IL, Perez-Quintero A, Booher NJ, Dossa GS, Grande G, Szurek B, Vera Cruz C, Bogdanove AJ, Oliva R. 2016. Effector diversification contributes to *Xanthomonas oryzae pv. oryzae* phenotypic adaptation in a semi-Isolated environment. Sci Rep 6:34137. doi:10.1038/srep34137

Richter A, Streubel J, Blücher C, Szurek B, Reschke M, Grau J, Boch J. 2014. A TAL effector repeat architecture for frameshift binding. Nat Commun 5:3447. doi:10.1038/ncomms4447

Sadoine M, Long J, Song C, Arra Y, Frommer WB, Yang B. 2021. Sucrose-dependence of sugar uptake, quorum sensing and virulence of the rice blight pathogen *Xanthomonas oryzae pv. oryzae*. doi: 10.1101/2021.08.22.457195

Schliep KP. 2011. phangorn: phylogenetic analysis in R. Bioinforma Oxf Engl 27:592–593. doi: 10.1093/bioinformatics/btq706

Séré Y, Onasanya A, Verdier V, Akator K, Ouedraogo LS, Segda Z, Coulibaly MM, Sido AY, Basso A. 2005. Rice bacterial leaf blight in West Africa: preliminary studies on disease in farmers’ fields and screening released varieties for resistance to the bacteria. Asian J Plant Sci 4:577–579. doi: 10.3923/ajps.2005.577.579

Streubel J, Pesce C, Hutin M, Koebnik R, Boch J, Szurek B. 2013. Five phylogenetically close rice SWEET genes confer TAL effector-mediated susceptibility to *Xanthomonas oryzae* pv. *oryzae*. New Phytol 200:808–19. doi: 10.1111/nph.12411

Tall H, Tékété C, Noba K, Koita O, Cunnac S, Hutin M, Szurek B, Verdier V. 2020. Confirmation report of bacterial blight caused by *Xanthomonas oryzae pv. oryzae* on rice in Senegal. Plant Dis 104:968–968. doi:10.1094/PDIS-07-19-1464-PDN

Tekete C, Cunnac S, Doucouré H, Dembele M, Keita I, Sarra S, Dagno K, Koita O, Verdier V. 2020. Characterization of new races of *Xanthomonas oryzae pv. oryzae* in Mali informs resistance gene deployment. Phytopathology 110:267–277. doi:10.1094/PHYTO-02-19-0070-R

Tran TT, Pérez-Quintero AL, Wonni I, Carpenter SCD, Yu Y, Wang L, Leach JE, Verdier V, Cunnac S, Bogdanove AJ, Koebnik R, Hutin M, Szurek B. 2018. Functional analysis of African *Xanthomonas oryzae pv. oryzae* TALomes reveals a new susceptibility gene in bacterial leaf blight of rice. PLoS Pathog 14:e1007092. doi:10.1371/journal.ppat.1007092

van der Tycho TA, Yonho K. 2022. On the success and failure of North Korean development aid in Africa. In: Yonho K, editor. NKEF Policy and Research Paper Series. Washington: George Washington University. pp. 31–42.

Vera Cruz CM, Bai J, Ona I, Leung H, Nelson RJ, Mew TW, Leach JE. 2000. Predicting durability of a disease resistance gene based on an assessment of the fitness loss and epidemiological consequences of avirulence gene mutation. Proc Natl Acad Sci U S A 97:13500–5. doi:10.1073/pnas.250271997

Verdier, Triplett LR, Hummel AW, Corral R, Cernadas RA, Schmidt CL, Bogdanove AJ, Leach JE. 2012a. Transcription activator-like (TAL) effectors targeting *OsSWEET* genes enhance virulence on diverse rice (*Oryza sativa*) varieties when expressed individually in a TAL effector-deficient strain of *Xanthomonas oryzae*. New Phytol 196:1197–1207. doi:10.1111/j.1469-8137.2012.04367.x

Verdier, Vera Cruz C, Leach JE. 2012b. Controlling rice bacterial blight in Africa: Needs and prospects. J Biotechnol 159:320–328.

Wick RR, Holt KE. 2022. Polypolish: Short-read polishing of long-read bacterial genome assemblies. PLoS Comput Biol 18:e1009802. doi:10.1371/journal.pcbi.1009802

Wu L-B, Eom J-S, Isoda R, Li C, Char SN, Luo D, Schepler-Luu V, Nakamura M, Yang B, Frommer WB. 2022. OsSWEET11b, a potential sixth leaf blight susceptibility gene involved in sugar transport-dependent male fertility. New Phytol 234:975–989. doi: 10.1111/nph.18054

Xu Z, Xu X, Gong Q, Li Z, Li Y, Wang S, Yang Y, Ma W, Liu L, Zhu B, Zou L, Chen G. 2019. Engineering broad-spectrum bacterial blight resistance by simultaneously disrupting variable TALE-binding elements of multiple susceptibility genes in rice. Mol Plant 12:1434–1446. doi:10.1016/j.molp.2019.08.006

Zetsche B, Gootenberg JS, Abudayyeh OO, Slaymaker IM, Makarova KS, Essletzbichler P, Volz SE, Joung J, van der Oost J, Regev A, Koonin EV, Zhang F. 2015. Cpf1 is a single RNA-guided endonuclease of a class 2 CRISPR-Cas system. Cell 163:759–771. doi:10.1016/j.cell.2015.09.038

Zhang Hui, Zhang J, Wei P, Zhang B, Gou F, Feng Z, Mao Y, Yang L, Zhang Heng, Xu N, Zhu J-K. 2014. The CRISPR/Cas9 system produces specific and homozygous targeted gene editing in rice in one generation. Plant Biotechnol J 12:797–807. doi:10.1111/pbi.12200

Zhou H, Liu B, Weeks DP, Spalding MH, Yang B. 2014. Large chromosomal deletions and heritable small genetic changes induced by CRISPR/Cas9 in rice. Nucleic Acids Res 42:10903–10914. doi: 10.1093/nar/gku806

